# From reactive to anticipatory control of tongue movements for interception in mice

**DOI:** 10.64898/2026.01.25.701632

**Authors:** Mohamed ElTabbal, Cedric Galetzka, Bernd Kuhn

**Author notes:** These authors contributed equally (Mohamed ElTabbal, Cedric Galetzka).

## Abstract

Object interception requires integrating prediction and sensory feedback, making it a powerful model for studying sensorimotor transformations. However, traditional head-fixed mouse paradigms reduce behavior to stereotyped stimulus–response associations and lack key features of natural interception. To address this limitation, we developed a behavioral task in which head-fixed mice use their tongue to intercept a food pellet moving at one of seven constant speeds randomly selected on each trial. We characterized the 3D kinematics of discrete tongue-reaching movements, their distinct motor phases, and learning-dependent changes in movement timing and kinematics. Mice adapted both tongue kinematics and movement onset to pellet speed, initially relying on reactive control and later adopting an anticipatory strategy. Anticipatory control depended largely on vision and intact lateral cerebellar circuits. This paradigm provides a powerful platform for investigating the neural mechanisms underlying predictive sensorimotor control and object interception in head-fixed mice.

**Significance Statement:** In natural environments, interception requires seamless sensory-motor transformation, yet traditional paradigms like classical conditioning and virtual reality fall short of capturing this complex behavior. We introduce a behavioral paradigm in which trained mice intercept food pellets moving at constant speeds that vary randomly across trials, with discrete tongue licks. Remarkably, mice exhibit a shift from a reactive to a more anticipatory motor strategy, adaptively tuning lick initiation and projection speed in a learning-dependent manner. This work establishes a simplified interception-like framework for studying sensorimotor dynamics in head-fixed mice.

## INTRODUCTION

Interception is an essential behavior observed across a broad spectrum of species in their natural environments, highlighting its utility as a paradigm for investigating sensory-motor transformation. This process involves continuously mapping sensory inputs into spatially and temporally coordinated motor commands. For example, dragonflies predictively orient their heads to track prey angular position, adjusting flight paths via rapid visual feedback^1^; salamanders extrapolate prey trajectories to project their tongues, mitigating sensorimotor delays^2^; and marmosets integrate stalking, gaze tracking, and ballistic grasping to capture targets^3^.

To dissect the process of sensory-motor transformation during interception under controlled conditions, researchers have developed behavioral paradigms for humans and monkeys, employing manipulanda or joysticks to guide a cursor on a two-dimensional plane^4,5^, or using rods or arms to strike moving objects^6^. These studies have yielded critical behavioral insights, such as coupling between object and hand velocities^4,6^, reduced reaction times for faster-moving targets, and the generation of complex velocity profiles including corrective submovements for slower targets^4^. In these tasks, interception relies on a mix of reactive and predictive strategies^4^, with visual input central to the latter^7^. Neuroimaging and lesion studies, predominantly in primates, have implicated various brain regions in the process of sensory-motor transformation during interception^8^.

While these findings illuminate behavioral and neural mechanisms underlying sensorimotor processing during interception, a comprehensive understanding of this dynamic sensorimotor process demands a genetically tractable mammalian model. Yet, interception-focused behavioral research in mice remains limited. Existing behavioral paradigms in mice typically emphasize either motor execution, such as forelimb reaching to static targets^9,10^, or sensory processing, such as responses to looming or moving visual stimuli^11,12^. A more naturalistic approach to studying interception behavior in freely moving mice as they adeptly intercept prey was previously reported^13,14^. Still, this task spans a vast parameter space of sensory and motor variables, making it more difficult for experimental control. To our knowledge, there is no behavioral paradigm in head-fixed mice investigating sensorimotor control using interception as a behavioral model.

In forelimb-reaching protocols, it was shown that, early in training, mice naturally retrieve static pellets with their tongues when within range^15,16^. Leveraging this tongue-reaching capability, we designed an interception behavior paradigm in head-fixed mice, tasking them with intercepting a pellet moving at constant speeds that change randomly from trial to trial using their tongue. This trial-to-trial randomization minimized reliance on memorized temporal intervals, encouraging mice instead to use ongoing sensory information about pellet motion to guide interception. We describe the construction of the experimental setup hardware and behavioral analysis.

Our results demonstrate that mice execute discrete tongue-reaching movements, adapting the onset of the tongue-reach movement (anticipatory strategy) or the kinematics of these movements (reactive strategy) in relation to the pellet speed to achieve successful interception. Additionally, we show that most mice employ an anticipatory strategy later in learning versus a more reactive one early in learning.

## 1 RESULTS

### 1.1 Experimental Setup and Tongue Tracking

The experimental setup consists of three primary components: a linear belt actuator rail and motor unit, an automatic pellet refilling mechanism, and two 500 Hz infrared cameras. These components are controlled by an Arduino microcontroller (see Methods for more information). This arrangement allowed us to automatically vary the speed of the moving platform on every trial, automatically refilling pellets between trials, and triggering simultaneous data acquisition (Figure 1A). Food pellets were placed on a platform connected to a micromanipulator, and on each trial, the food pellet moved at a randomly selected speed from 1 to 7 cm/s (step size: 1 cm/s). Following handling and habituation, mice completed up to six sessions (one session per day) in which they learned to intercept the moving food pellets with their tongue (Figure S2A). We extracted whole-body motion and whisker pad activity from the high-speed video recordings during the first and last sessions (Figure 1B). In addition, we tracked the tongue using DeepLabCut^17^ and analyzed tongue trajectories and kinematics in detail (Figure 1C).

**Figure 1:**
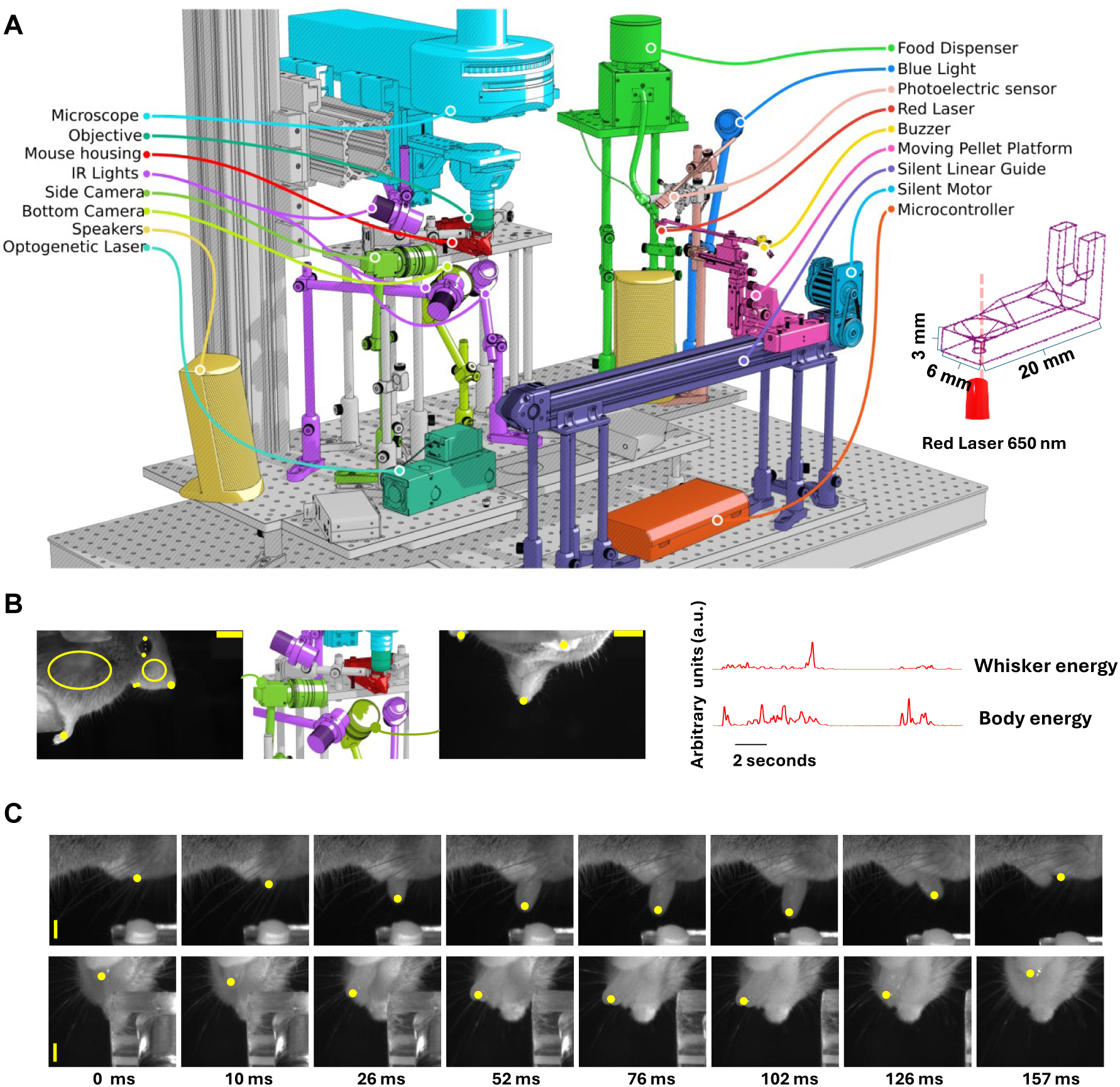
Schematic of the experimental setup and its components. (A) Experimental setup. All components except the microscope are mounted on aluminum breadboards. The system is controlled by an Arduino microcontroller and can be integrated with microscopy and optogenetics. The subpanel shows the design of the pellet platform. If no food pellet is present, a red laser emits laser light through the pellet platform, which a photo-sensor detects, triggering pellet release from the food dispenser. (B) Left: Sideand bottom-view cameras. Yellow dots indicate tracked points near the eyelid, snout, right paw, and upper and lower jaw using DeepLabCut^17^. Open yellow circles denote regions of interest used to analyze the motion energy of the whisker pad and body. Middle: The high-speed cameras and surrounding LEDs. Right: Example traces generated by the motion energy analysis module. All traces were normalized. Scale bar = 6 mm. (C) Example of tongue tip tracking using DeepLabCut^17^ and further refinement via the manual tongue tracking module. The lick lasted for 157 milliseconds. Scale bar = 3 mm.

To analyze tongue trajectories, we segmented each lick into projection, probing, and retraction phases (Figures S1A, S1B; see Methods for more information). We noticed that mice performed one of two distinct lick types: the first type, simple licks, followed a simple progression of these motor phases with variable timing of the probing phase (Figure S1B), whereas the second type, complex licks, exhibited multiple transitions between motor phases (Figure S1D; Supplementary movie 1). These transitions represent repeated interception attempts and can, therefore, be classified as online movement corrections. The online corrective movements of the mouse tongue resemble those of the human hand when reaching for a moving target^18^, demonstrating similar adaptive control and feedback-based adjustments. However, most licks were simple licks (mean = 93.7%± 0.97; SEM; Figure S1C), and the percentage of complex licks stayed low, with no significant change after learning (Figure S2D). We analyzed different lick parameters across all trials, regardless of lick type. These parameters included lick onset distance, defined as the distance between the midpoint of the mouse and the right edge of the food pellet platform at the time of lick onset, lick projection speed, lick duration, and lick endpoint, defined as the 3D location of the tongue tip at the onset of the last retraction phase.

### 1.2 Behavioral Performance and Motor Learning

We analyzed learning-dependent changes in trial success, lick distance onset, fine-scale kinematics, and body and whisker movements. To assess general performance measures, we classified trials as ‘hits’ if a mouse intercepted the pellet and raised it more than 2 mm using its tongue. Trials with licks that did not contact the pellet were classified as ‘misses’, and trials without licks as ‘no-lick’ trials. We compared the first (early) and last (late) training sessions. Mice in the late session showed a significant increase in the hit rate, and this increase was pronounced at pellet speeds higher than 3 cm/s (Figure 2A, left panel). The hit rate evolved gradually over the six training sessions and progressed more slowly for higher pellet speeds (Figure S2A, middle and right panels). The miss rate showed the opposite pattern, and the no-lick rate decreased with learning, especially at pellet speeds higher than 5 cm/s (Figure 2A, middle and right panels). Since trials with different pellet speeds were randomly interleaved, no-lick trials reflect trial-by-trial variability in performance, as mice frequently initiated licking or successfully intercepted the pellet on subsequent trials with different speeds.

**Figure 2:**
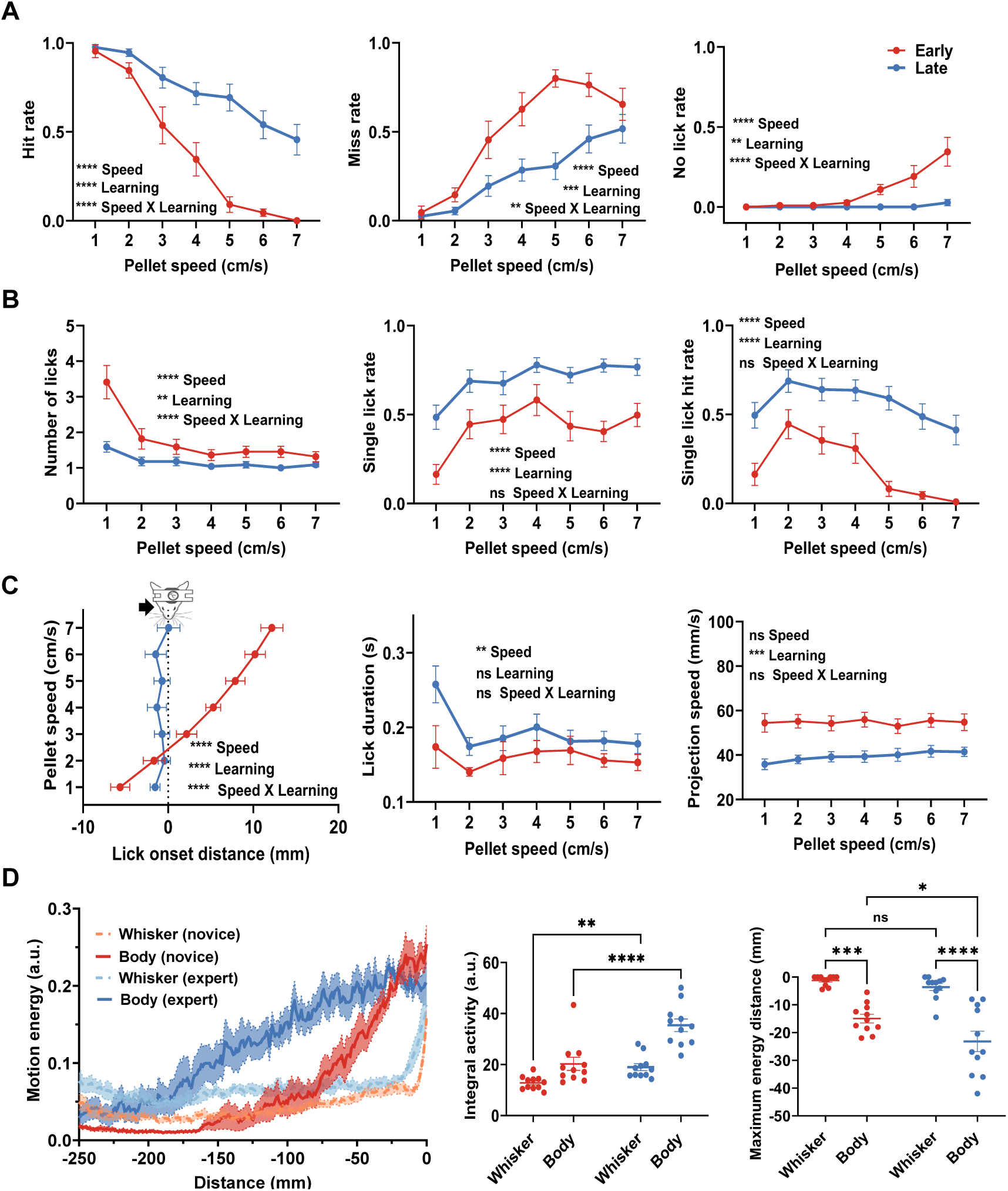
Quantitative measures of behavioral performance and motor learning. (A) Left: Hit rate for early versus late session mice. A two-way repeated measures ANOVA revealed significant main effects of learning (*F* (1, 10) = 69.06*, p <* 0.0001) and pellet speed (*F* (6, 60) = 57.71*, p <* 0.0001). The interaction between learning and pellet speed was significant as well (*F* (6, 60) = 9.762*, p <* 0.0001). Middle: Same as the left panel, but for the missing rate. There was a significant main effect for learning (*F* (6, 60) = 28.32*, p <* 0.0001) and pellet speed (*F* (1, 10) = 31.45*, p* = 0.0002) as well as their interaction (*F* (6, 60) = 3.809*, p* = 0.002). (A) Right: Same as the left panel, but for the no-lick rate. There was a significant main effect for learning (*F* (1, 10) = 12.93*, p* = 0.005), pellet speed (*F* (6, 60) = 11.33*, p <* 0.0001), and their interaction effect (*F* (6, 60) = 10.71*, p <* 0.0001). (B) Left: Number of licks per trial for early versus late session mice. A two-way repeated measures ANOVA revealed significant main effects for learning (*F* (1, 10) = 10.61*, p* = 0.0086) and pellet speed *F* (6, 60) = 17.53*, p <* 0.0001). The interaction between learning and pellet speed was significant as well (*F* (6, 60) = 8.605*, p <* 0.0001). Middle: Same as the left panel, but for the singlelick rate. There was a significant main effect for learning (*F* (1, 10) = 45.67*, p <* 0.0001) and pellet speed (*F* (6, 60) = 6.76*, p <* 0.0001), but the interaction was not significant (*F* (6, 60) = 0.775*, p* = 0.5921). Right: Same as the left panel, but for the single-lick hit rate. There was a significant main effect for learning (*F* (1, 10) = 82.05*, p <* 0.0001), pellet speed (*F* (6, 60) = 6.622*, p <* 0.0001), and a non-significant interaction effect (*F* (6, 60) = 1.568*, p* = 0.1723). (C) Left: Lick onset distance as a function of pellet speed. A two-way repeated measures ANOVA showed a significant main effect for learning (*F* (1, 10) = 61.59*, p <* 0.0001) and pellet speed (*F* (6, 60) = 46.68*, p <* 0.0001) as well as a significant interaction (*F* (6, 60) = 32.18*, p <* 0.0001). Middle: Same as the left panel, but for lick duration. There was a significant main effect for learning (*F* (1, 10) = 4.46*, p* = 0.0607) and pellet speed (*F* (6, 60) = 3.209*, p* = 0.0085) and a non-significant interaction (*F* (6, 60) = 1.746*, p* = 0.1260). Post-hoc Bonferroni comparisons revealed that the slowest pellet speed was associated with the longest lick duration. Right: Same as in the left panel, but for lick projection speed. There was a significant main effect for learning (*F* (1, 10) = 24.88*, p* = 0.0005) but a nonsignificant main effect of pellet speed *F* (6, 60) = 0.7649*, p* = 0.6004 and interaction (*F* (6, 60) = 0.9102*, p* = 0.4939). (D) Left: Mean whole-body and whisker motion energy in early and late session mice as a function of pellet distance from the midpoint of the mice. Middle: Integral of whole-body and whisker motion energy across distance in early and late session mice. A two-way repeated measures ANOVA followed by multiple comparisons using the Bonferroni correction revealed significant differences between early and late session mice for the integral motion of the body (p*<*0.0001) and whiskers (p*<*0.0040). Right: Average distance between the pellet and the midpoint of the mice at peak whole-body and whisker motion energy in early and late session mice. A two-way repeated measures ANOVA followed by multiple comparisons using Bonferroni correction revealed significant differences between early and late session mice for the distance of maximum activity of the body (mean difference = 8.22 mm, p*<*0.0001) but not of the whiskers (p*>*0.05). The distance at which body movement reached its peak preceded whiskers in mice in the early (mean difference=13.73 mm, p=0.0008) and late sessions (mean difference = 19.59 mm, p*<*0.0001). Error bars represent the SEM (N = 11 mice).

To evaluate the contribution of non-rhythmic licking in this task, we analyzed trials with single, discrete licks. As mice learned, the number of licks per trial decreased, and this decrease was more pronounced at slower speeds (Figure 2B, left panel). There was a significant increase in the single-lick rate at different pellet speeds (Figure 2B, middle panel). Additionally, the accuracy of these single-licks increased significantly later in learning, as shown by the increase in the single-lick success rate (Figure 2B, right panel). Figure S4 illustrates an example of a single mouse tongue-reaching behavior and how it evolves from rhythmic licking to more discrete, single tongue-reaching movements. These findings suggest that mice learn to initiate non-rhythmic, discrete single licks to successfully intercept the food pellet.

In interception tasks, the critical variables are the position and speed of the moving object, as successful interception requires a well-timed motor response that accounts for both. To capture the spatial context of the animal’s response, we analyzed lick-onset distance (the distance between the right edge of the pellet platform and the mouse’s midline at the onset of tongue movement). This measure provides direct information about the pellet’s position in space when movement is initiated and, when considered alongside pellet speed, offers insight into how the animal coordinates its action in response to varying pellet speeds. We also analyzed other kinematic parameters (e.g., lick projection speed and lick duration) at the early and late learning stages to understand learning-related changes in the fine-scale parameters of tongue-reaching movements.

In early learning, at slower pellet speeds (1 and 2 cm/s), mice initiated licks at a greater distance before the pellet reached them. In comparison, at higher pellet speeds (3-7 cm/s), mice initiated their licks after the pellet passed them (Figure 2C, left panel). With learning, mice shifted their first lick onset distance in all speeds to slightly precede the food pellet’s arrival at the midpoint (Figure 2C, left panel). Initially, this suggested that mice in late sessions learn to lick at a fixed distance (a threshold-distance strategy), rather than adjusting for pellet speed (starting at earlier distances for faster pellet speeds). This latter pattern would be expected if the mouse is estimating the pellet’s arrival time at the midline. We reasoned that if the threshold-distance strategy was used for successful interception in late sessions, successful first licks would exhibit the same trend. However, when looking into the first successful licks, we found that mice in late sessions adjusted their lick onset distance based on pellet speed, initiating their licks at earlier distances for higher pellet speeds (Figure S2B, left panel). More importantly, this first successful lick pattern is not found in mice in early sessions, where lick onset distances have a slight positive deviation from the midline (Figure S2B, left panel), indicating that, as mice become more trained, they adaptively adjust their lick onset distance in an anticipatory, learning-dependent manner during successful interceptions.

First lick duration showed an insignificant increase between early and late learning sessions. The lowest pellet speed was associated with the longest lick duration, but there was no pellet speed-dependent effect over learning (Figure 2C, middle panel). We observed the same pattern for successful first licks (Figure S2B, middle panel). In addition, the mean projection speed of all first licks showed a significant decrease in mice with no pellet speed-dependent effects (Figure 2C, right panel), though successful first licks in the early session tended to be faster for higher pellet speeds (Figure S2B, right panel). The spatial endpoint of licks did not show a significant difference in the mediolateral, dorsoventral, or anteroposterior axes (Figure S2C). These results indicate that all mice adaptively adjusted their lick onset distance and projection speed during learning (see Supplementary Movie 2). Additionally, mice seem to fine-tune these parameters in response to variations in pellet speed. In contrast, lick duration and the endpoint’s spatial coordinates showed no learning-dependent changes.

To investigate preparatory body movement during the task, we analyzed whole-body and whisker motion energy traces as the food pellet approached the mice from approximately 250 mm relative to the midline. Whole-body and whisker movements exhibited earlier ramping in mice in the late session compared to the early training session (Figure 2D, left panel). There was an increase in whole-body and whisker movements in mice in the late session compared to the early session (Figure 2D, middle panel). In addition, we measured the distance between the pellet and the midline of the mouse at the maximum movement intensity of either the whisker or the whole body. Mice in the late session shifted the peak of their whole-body movement earlier relative to the early training session, and whole-body movements consistently peaked earlier than whisker movements (Figure 2D, right panel). These results suggest that mice show coordinated preparatory body and whisker movements with learning.

### 1.3 Behavioral Strategies for Successful Interception

Behavioral strategies are valuable when interacting with a dynamic environment^19^. A main feature of the interception task was that pellet speed changed randomly from trial to trial, requiring animals to adapt their movement parameters on a trial-by-trial basis. We found that mice in the late session adjusted their lick onset distance in response to different pellet speeds to achieve successful interception. In contrast, mice in the early session scaled their tongue projection speed to successfully intercept pellets at higher speeds. This suggests that mice use two behavioral strategies to achieve successful interception.

We used a logistic linear mixed-effects model (LLMM) to determine which tongue kinematic and task factors predict successful pellet interception and how their interaction changes across pellet speeds and sessions. Fixed effects in the model included three lick metrics, lick onset distance (Lick O), lick duration (Lick D), and lick projection speed (Proj Sp), as well as pellet speed (Pel Sp), which has seven levels, and the animal’s learning stage (early vs. late session). To capture the dependencies between these predictors, we also added all possible two-way interactions (e.g., Lick O × Pel Sp, Lick D × Learning Stage) and the three-way interaction (e.g., Lick O × Pel Sp × Learning Stage). Mouse identity was treated as a random effect, allowing each animal to have its own intercept (baseline success probability) and slope with pellet speed. In this framework, the fixed-effect coefficients represent the change in the likelihood of successful interception at the reference condition (pellet speed = 1 cm/s, early learning), the two-way interaction terms show how one predictor’s effect varies across levels of another, and the three-way interaction captures any additional modulation of those pairwise effects.

Early in learning (Figure 3A), at the reference pellet speed (1 cm/s), none of the individual kinematic features, lick duration, onset distance, or tongue projection speed, significantly influenced the likelihood of successful interception. However, a significant interaction between tongue projection speed and pellet speed was observed (*β* = 0.516, *p* = 0.003), indicating that increasing the tongue’s projection speed became beneficial as pellet speed increased. At this early learning stage, pellet speed had a strong negative main effect on success (*β* = -1.377, *p <* 0.0001), suggesting that mice in the early session were less able to adjust to faster-moving targets. We interpret this pattern as a relatively reactive response (i.e., projecting faster when the target moves faster to compensate for a late start of the lick) rather than precise timing coordination.

**Figure 3:**
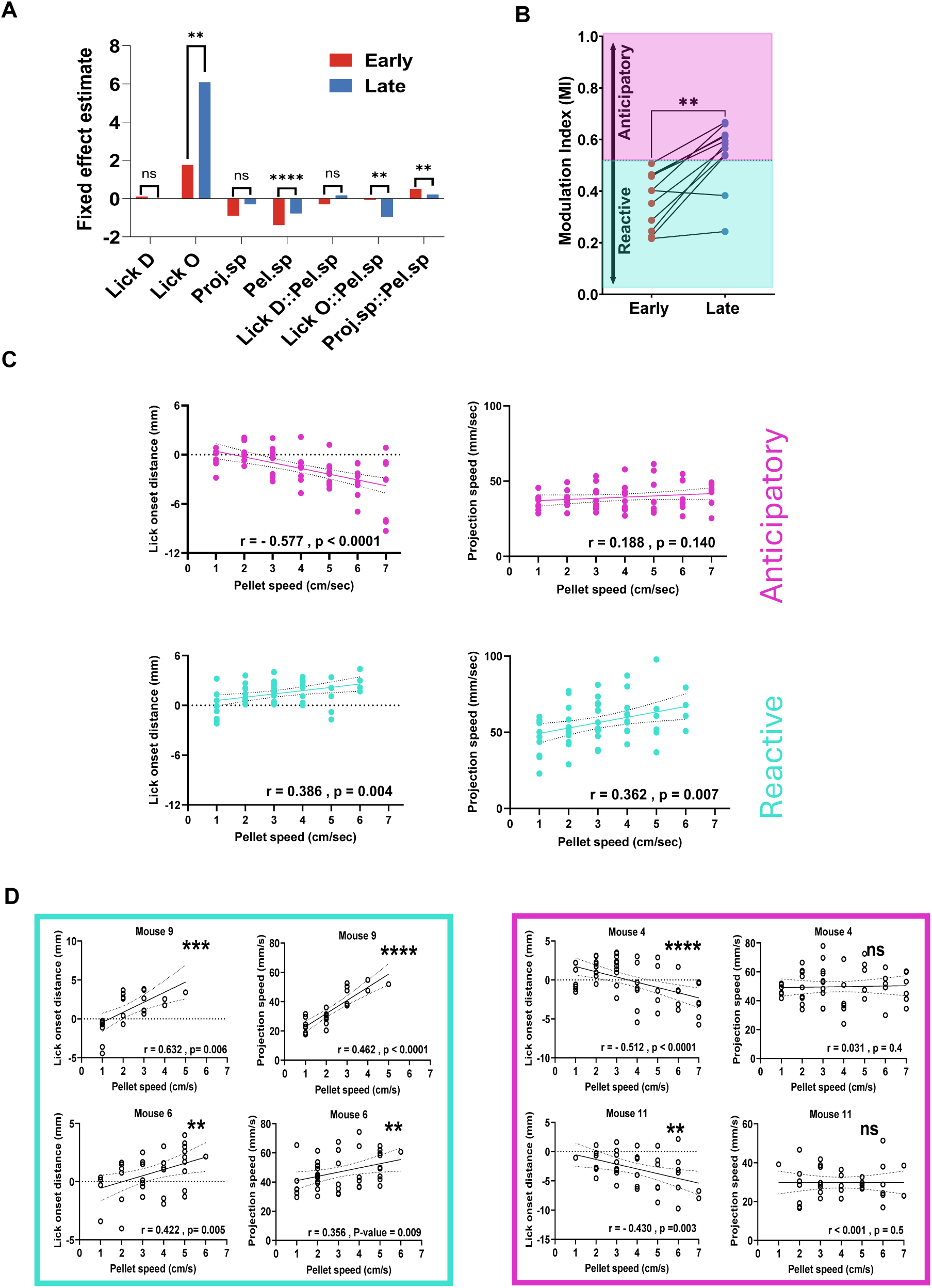
Behavioral strategies for interception. (A) Results of the logistic linear mixed effect model, quantifying the relationship between behavioral and task variables and the probability of successful interception. (A) A positive fixed effect estimate means higher chances of successful interception. Lick onset distance (Lick O), lick duration (Lick D), and lick projection speed (Proj.sp) are assessed at the reference level of pellet speed (Pel.sp) 1 cm/s. The interaction terms show how the reference level of the fixed effect is modulated by increasing pellet speed. Significant changes from the early session (red) to the late session (blue) are shown according to their level of significance (*p*<*0.05, **p*<*0.01, ***p*<*0.001, and ****p*<*0.0001) (B) Strategy modulation index (MI) for early and late session mice. Each dot represents a single mouse. The MI represents a continuum, with higher scores indicating a more anticipatory strategy and lower scores indicating a more reactive strategy. The dotted line represents a threshold (the mean of the centroids of two clusters from k-means clustering of all MI scores), dividing the more reactive (cyan shading) from the anticipatory mice (pink shading). A paired t-test revealed a significant difference in the MI between the early and late sessions (p = 0.002). (C) Lick onset distance or tongue projection speed versus pellet speed for the anticipatory and reactive clusters. Each dot represents the mean lick onset distance or lick projection speed value of a single mouse clustered in either of the two groups. (D) Left: Examples of two mice in the reactive cluster (cyan box). Right: Same as the left panel, but for mice in the anticipatory cluster (pink box). Pearson correlation coefficients are reported alongside p-values from two-tailed significance tests.

Later in learning (Figure 3A), the influence of pellet speed on success was significantly attenuated (Pellet Speed × Learning Stage: *β* = 0.602, *p <* 0.0001), reflecting improved behavioral adaptation. Crucially, lick onset distance emerged as a significant predictor of success at low pellet speeds in mice during the late session (Lick Onset distance × Learning Stage: *β* = 4.326, *p* = 0.007), suggesting that the timing of movement initiation had become more critical. However, this benefit was diminished at higher pellet speeds, as indicated by a significant negative three-way interaction between lick onset distance, pellet speed, and learning stage (*β* = -0.893, *p* = 0.002). This suggests that intercepting fast-moving pellets required earlier, more precisely timed licks, and that late licks became detrimental as pellet speed increased. Notably, the previously beneficial interaction between tongue projection speed and pellet speed became significantly reduced in this late learning stage (Projection Speed × Pellet Speed × Learning Stage: *β* = -0.302, *p* = 0.004), implying a shift away from relying on the more reactive pattern seen in earlier learning stages.

Based on the results of the LLMM, we extracted three behavioral features for each mouse: the correlation between lick onset distance and pellet speed, the correlation between tongue projection speed and pellet speed, and the median value of the lick onset distance across successful trials. These features were then combined into a single index, the strategy modulation index (MI), to quantify how mice fine-tune their lick onset distance and the dependencies of both lick onset distance and tongue projection speed on pellet speed (MI; for more information, see Methods). The MI represents a continuum of learning, ranging from zero (indicating a more reactive approach) to one (indicating a more anticipatory approach). Mice at the late training session significantly improved their MI score compared to the early training session (median MI score difference = 0.1603, p = 0.002) (Figure 3B). K-means clustering was subsequently applied to all the MI scores at different learning stages to identify potential clusters (Figure 3B).

All mice in the early session were clustered together. As mice transitioned to the late session, most were clustered together except for two mice who remained in the first cluster. To understand how these two clusters differ in their lick onset distance, tongue projection speed, and pellet speed correlations, we plotted how these correlations differ in successful attempts for each cluster. While the more anticipatory cluster showed a significant negative correlation between lick onset distance versus pellet speed and an insignificant correlation between tongue projection speed and pellet speed (Figure 3C, upper panel). The other cluster (more reactive) showed a different pattern of significant positive correlations for both lick onset distance and tongue projection speed with pellet speed (Figure 3C, lower panel). This analysis suggests that mice use different successful interception strategies at different learning stages. While all mice shifted their approach from a more reactive to a more anticipatory strategy, two of these mice at the later learning stage did not shift their reactive strategy, showing similar patterns to the early learning stage (Figure 3D)

### 1.4 The Role of Vision in Anticipatory Interception

We then examined whether mice in the late session used visual information for successful interception. We trained four mice for up to six sessions, recorded their baseline performance, and then blocked their left visual field with a barrier (Figure 4A). When vision was blocked, the hit rate decreased, and this decrease was pronounced for higher pellet speeds (Figure 4B, left panel). Similarly, the miss rate increased under visual blocking, while the no-lick rate only increased for higher pellet speeds (Figure 4B, middle and right panels). There was a general increase in the average number of licks under visual blocking (Figure 4C, left panel) with a corresponding decrease in the single-lick rate (Figure 4C, middle). Additionally, there was a decrease in single-lick success rate (Figure 4C, right panels), indicating reduced accuracy. This data suggests that mice benefit from visual information for successful interception.

**Figure 4:**
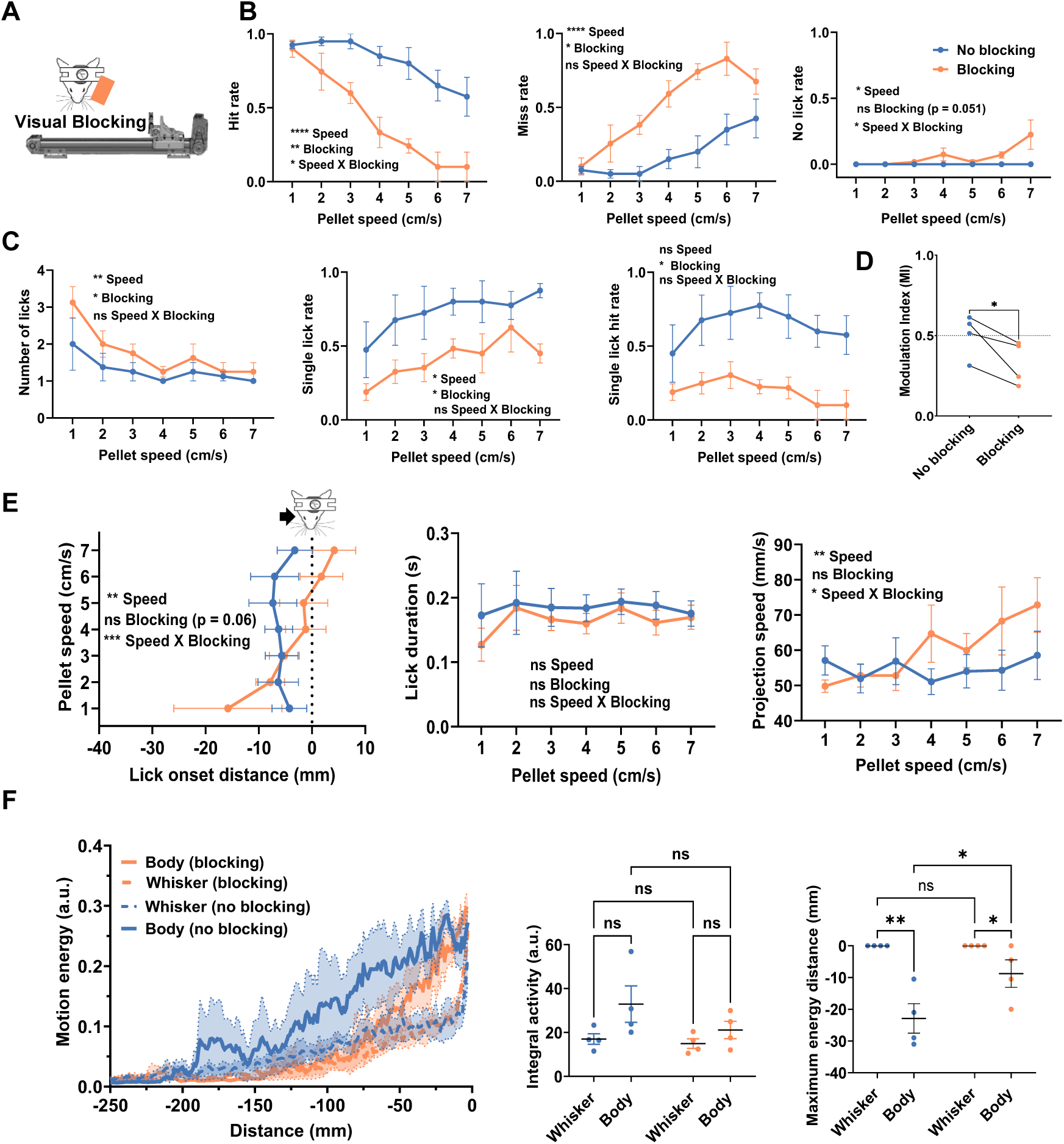
Effects of visual blocking on task performance, lick parameters, and body movement. (A) Schematic representation of the visual blocking experiment, in which the left visual field of mice in late learning sessions was blocked. (B) Left: Hit rate for all pellet speeds. A two-way repeated measures ANOVA revealed significant main effects for blocking (*F* (1, 3) = 48.20*, p* = 0.0061), pellet speed (*F* (6, 18) = 14.89*, p <* 0.0001), as well as their interaction (*F* (6, 18) = 3.116*, p* = 0.0284). Middle: Same as the left panel, but for the miss rate. There were significant main effects for blocking (*F* (1, 3) = 28.04*, p* = 0.0131) and pellet speed (*F* (6, 18) = 14.74*, p <* 0.0001), but the interaction was not significant (*F* (6, 18) = 1.999*, p* = 0.1191). (B) Right: Same as the left panel, but for the no-lick rate. There was a significant main effect for pellet speed (*F* (6, 18) = 2.905*, p* = 0.0368), a non-significant main effect for blocking (*F* (1, 3) = 9.249*, p* = 0.0558), and a significant interaction (*F* (6, 18) = 2.905*, p* = 0.0368). (C) Left: Number of licks in trials with at least one lick for all pellet speeds. A two-way repeated measures ANOVA revealed significant main effects for blocking (*F* (1, 3) = 11.27*, p* = 0.0438), pellet speed (*F* (6, 18) = 5.463*, p* = 0.0023), and a non-significant interaction (*F* (6, 18) = 0.8906*, p* = 0.5221). Middle: Same as the left panel, but for the single-lick rate. There were significant main effects for blocking (*F* (1, 3) = 11.41*, p* = 0.0432) and pellet speed (*F* (6, 18) = 3.335*, p* = 0.0218) and a non-significant interaction (*F* (6, 18) = 0.4498*, p* = 0.8358). Right: Same as the left panel, but for the single-lick hit rate. The main effect for blocking was significant (*F* (1, 3) = 25.77*, p* = 0.0148), but the main effect for pellet speed (*F* (6, 18) = 0.8372*, p* = 0.5572) and their interaction (*F* (6, 18) = 0.6525*, p* = 0.6880) were not significant. (D) Strategy modulation index for all mice during visual blocking and no blocking. A paired t-test showed a significant difference between blocking and no blocking conditions (mean difference = -0.1736, p=0.0494). (E) Left: Lick onset distance for all pellet speeds. A two-way repeated measures ANOVA showed a significant main effect for pellet speed (*F* (6, 18) = 6.592*, p* = 0.0008), a non significant main effect for blocking (*F* (1, 3) = 9.678*, p* = 0.0629), and a significant interaction (*F* (6, 18) = 6.929*, p* = 0.0006). Middle: Same as the left panel, but for lick duration. There were no significant main effects or interactions (all p *>* 0.05). Right: Same as the left panel, but for the lick projection speed. There was a significant main effect for pellet speed (*F* (6, 18) = 4.020*, p* = 0.009), a non-significant main effect for blocking (*F* (1, 3) = 2.530*, p* = 0.2096), and a significant interaction (*F* (6, 18) = 3.289*, p* = 0.0230). (F) Left: Mean whole-body and whisker motion energy during visual blocking and no blocking as a function of pellet distance from the midpoint of the mice. Middle: Integral of whole-body and whisker motion energy across distance. A two-way repeated measures ANOVA followed by multiple comparisons using Bonferroni correction revealed non-significant differences across all comparisons (all p *>* 0.05). Right: A two-way repeated measures ANOVA followed by multiple comparisons using Bonferroni correction revealed a significant difference (p*<*0.0001) in the distance at which body motion peaked before visual blocking (mean distance = -22.875 ± 4.656 mm) and after blocking (mean distance = -8.750 ± 4.323 mm), but not for whisker motion (p*>*0.05), which consistently reached its maximum activity when the pellet reached the midline of the animal. Error bars represent the SEM across mice in all panels (N = 4 mice).

Since mice in their late training session employ a more anticipatory strategy, licking at earlier distances for faster pellet speeds, we hypothesized that blocking visual information would bias them toward a more reactive strategy. As expected, the MI decreased during visual blocking for all trained mice in the late training session (Figure 4D). After visual blocking, lower pellet speeds were associated with earlier lick distance onset and higher pellet speeds with delayed lick distance onset (Figure 4E, left panel). Lick duration was not altered during visual blocking (Figure 4E, middle panel). However, even though visual blocking did not lead to a general increase in tongue projection speed, mice increased their tongue projection speed for higher pellet speeds (Figure 4E, right panel). These findings suggest that visual blocking leads to a delay in the lick onset distance when intercepting higher pellet speeds, which mice compensate for by increasing tongue projection speed. Hence, mice shifted towards a more reactive interception strategy when visual input was blocked.

We analyzed whole-body and whisker motion to examine the effect of visual blocking on preparatory movements (Figure 4F, left panel). Overall, body and whisker activity showed an insignificant decrease with a later onset of ramping (Figure 4F, middle panel). However, the distance at which mice exhibited maximum whole-body movement shifted closer to the midpoint of the mice. Maximum whisker movement timing remained unaffected by visual blocking (Figure 4F, right panel). These results show that the timing of preparatory body movements was altered under visual blocking.

### 1.5 Lateral Cerebellar Circuits’ Role in Anticipatory Interception

Since lateral cerebellar circuits have been shown to play an important role in anticipatory action in tasks similar to our interception paradigm across different model organisms^20,21^, we pharmacologically inactivated cerebellar cortical lobules Crus I and II. After six days of training, we recorded performance in four mice after unilateral cerebellar cortical injections of saline or muscimol, 150 µm below the surface (Figure 5A). Muscimol injections were associated with a decrease in the hit rate and an increase in the miss rate across all pellet speeds (Figure 5B, left and middle panel). There was no general increase in the rate of no-lick trials, though muscimol injection led to an increase in the no-lick rate at higher pellet speeds (Figure 5B, right panel). Furthermore, inactivating Crus I and II tended to increase the number of licks per trial for slower pellet speeds (Figure 5C, left panel). While the single-lick rate did not show a general change between saline and muscimol injections, the single-lick hit rate decreased across all pellet speeds. Therefore, inactivating lateral cerebellar circuits degrades task performance and the accuracy of single, discrete licks.

**Figure 5:**
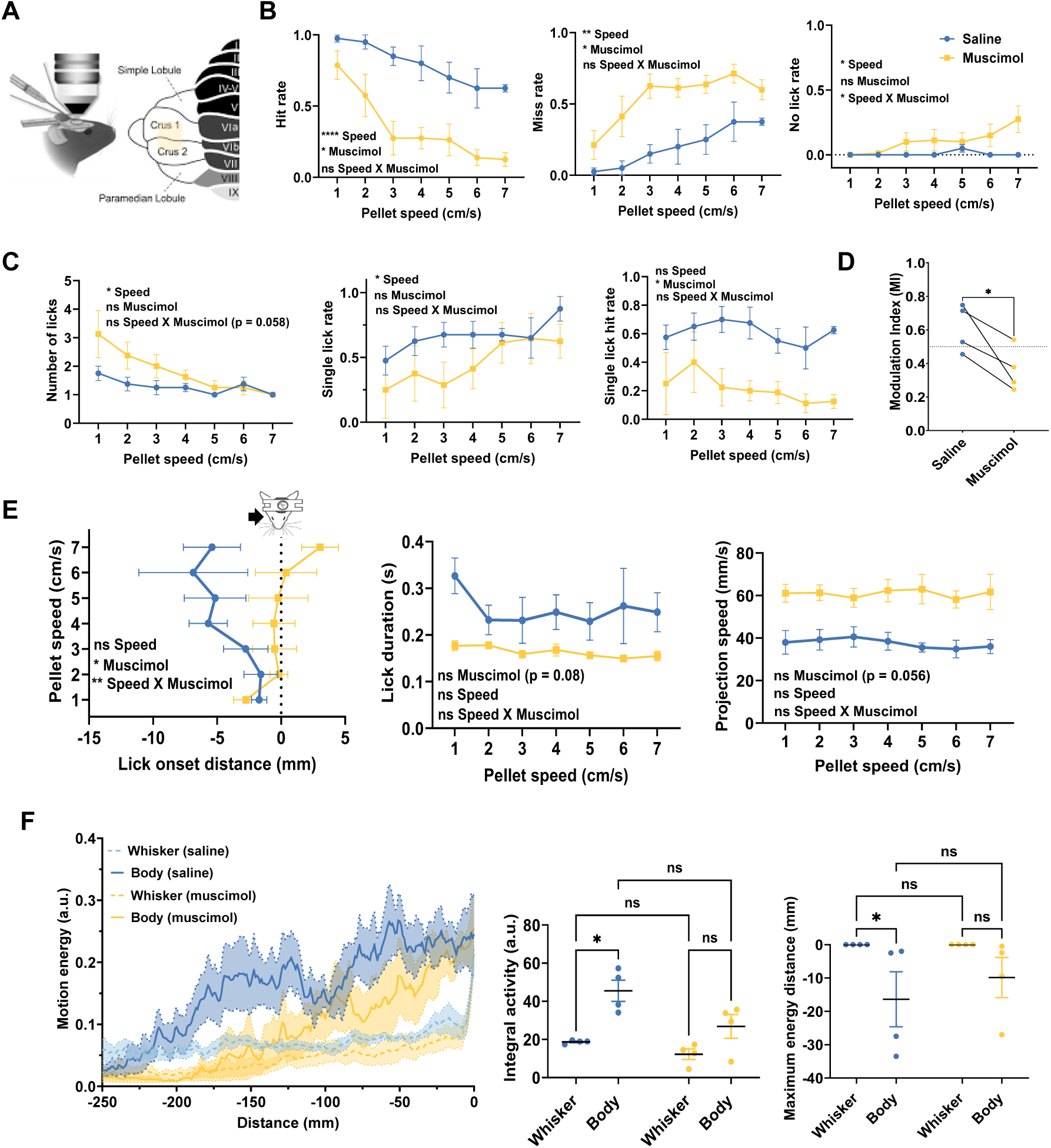
Effects of muscimol on task performance, lick parameters, and body movement. (A) Schematic representation of the intracranial saline and muscimol injections in mice in their late session. (B) Left: Hit rate for all pellet speeds. A two-way repeated measures ANOVA revealed significant main effects for muscimol (*F* (1, 3) = 17.12*, p* = 0.0256), pellet speed (*F* (6, 18) = 12.84*, p <* 0.0001), but not their interaction (*F* (6, 18) = 1.473*, p* = 0.247). Middle: Same as the left panel, but for the miss rate. There were significant main effects for muscimol *F* (1, 3) = 19.04*, p* = 0.0222) and pellet speed (*F* (6, 18) = 6.050*, p* = 0.0013), but the interaction was not significant (*F* (6, 18) = 0.7593*, p* = 0.7593). (B) Right: Same as the left panel, but for the no-lick rate. There was a significant main effect for pellet speed (*F* (6, 18) = 3.364*, p* = 0.0211), a non-significant main effect for muscimol (*F* (1, 3) = 2.911*, p* = 0.1863), and a significant interaction (*F* (6, 18) = 3.942*, p* = 0.0108). (C) Left: Number of licks in trials with at least one lick for all pellet speeds. A two-way repeated measures ANOVA revealed a non-significant main effect for muscimol (*F* (1, 3) = 4.899*, p* = 0.1137), a significant main effect for pellet speed (*F* (6, 18) = 5.357*, p* = 0.0025), and a non-significant interaction (*F* (6, 18) = 2.544*, p* = 0.0581). The post hoc analysis showed significant effects at low pellet speeds less than 4 cm/s.Middle: Same as the left panel, but for the single-lick rate. There was a non-significant main effect for Muscimol (*F* (1, 3) = 2.893*, p* = 0.1875), a significant main effect for pellet speed (*F* (6, 18) = 2.901*, p* = 0.0370), and a non-significant interaction (*F* (6, 18) = 1.343*, p* = 0.289). The post hoc analysis showed the significant reduction in rate was mainly in pellet speeds 2 cm/s, 3 cm/s, 4 cm/s, and 7 cm/s. Right: Same as the left panel, but for the single-lick hit rate. The main effect for Muscimol was significant (*F* (1, 3) = 15.03*, p* = 0.0304), but the main effect for Speed (*F* (6, 18) = 0.787*, p* = 0.591) and their interaction (*F* (6, 18) = 0.9344*, p* = 0.494) were not significant. (D) Strategy modulation index (MI) for all mice following Muscimol or saline injection. A paired t-test showed a significant difference between Muscimol and saline (mean difference = -0.2484, p=0.0402). (E) Left: Lick onset distance for all pellet speeds. A twoway repeated measures ANOVA showed a non-significant main effect for pellet speed (*F* (6, 18) = 0.8107*, p* = 0.5751), a significant main effect for Muscimol (*F* (1, 3) = 10.84*, p* = 0.046), and a significant interaction (*F* (6, 18) = 5.791*, p* = 0.0017). Post hoc analysis shows a significant effect at all high pellet speeds *>* 4cm/sec. Middle: Same as the left panel, but for lick duration. There was no significant main effect for Muscimol (*F* (1, 3) = 6.562*, p* = 0.083), a significant main effect for pellet speed (*F* (6, 18) = 1.121*, p* = 0.3889), and a non-significant interaction (*F* (6, 18) = 0.8661*, p* = 0.538). Right: Same as the left panel, but for the lick projection speed. There was not significant main effect for Muscimol (*F* (1, 3) = 9.131*, p* = 0.0567) but with very large effect size. Pellet seed effect was not significant (*F* (6, 18) = 0.5474*, p* = 0.7656), same for the interaction between pellet speed and Muscimol effects (*F* (6, 18) = 0.8162*, p* = 0.5714). (E) Left: Mean traces of whole-body and whisker motion energy after muscimol or saline injection as a function of pellet distance from the midpoint of the mice. Middle: Integral of the left panel’s whole-body and whisker motion energy traces. A two-way repeated measures ANOVA followed by multiple comparisons using Bonferroni’s correction revealed a non-significant difference between body and whisker movement in saline and Muscimol conditions (p *>* 0.05). Right: A two-way repeated measures ANOVA followed by multiple comparisons using Bonferroni’s correction revealed a non-significant difference in the distance at which body motion peaks in saline and Muscimol conditions. The same goes for the whisker (p*>*0.05), which reached maximum activity consistently when the object reached the animal’s midline. Error bars represent SEM across mice in all panels (N = 4 mice).

We then analyzed lick parameters to examine whether unilateral muscimol injection led to a shift from anticipatory to more reactive behavior. The MI showed a decrease following muscimol compared to saline injection in all mice (Figure 5D). In addition, muscimol injection caused a general delay in lick onset distance pronounced at pellet speeds 4-7 cm/s (Figure 5E, left panel). Muscimol also tended to decrease lick duration, but this effect was not significant (Figure 5E, middle panel). Additionally, muscimol increased tongue projection speed across all pellet speeds (Figure 5E, right panel). We then analyzed whether this unilateral muscimol injection biased the lick angle. There was an increase in the lick angle ipsilateral to the muscimol injection site compared to saline. Additionally, the distribution of lick angles after muscimol injection became broader (Figure S3A). To further confirm this result, we compared the lick endpoint in all three axes (anteroposterior, dorsoventral, and mediolateral). Only the mediolateral axis exhibited significant lateralization of lick endpoints ipsilateral to the injection site (Figure S3B). These effects show that the inactivation of lateral cerebellar circuits not only impairs anticipatory behavior by delaying lick onset distances at higher pellet speeds but also changes the kinematics of this discrete tongue-reaching movement.

To understand if muscimol also affected whisker and body movement, we compared the motion energy traces following saline and muscimol injection (Figure 5F, left panel). There was no significant decrease in general body and whisker movement after muscimol compared to saline injection (Figure 5F, middle panel). In addition, the distance at which mice exhibited maximum whole-body movement slightly shifted towards the midpoint of the mice after muscimol compared to saline injection, but this effect was not significant (Figure 5F, right panel). Overall, muscimol did not significantly impair whole-body and whisker-coordinated movements.

## 2 Discussion

We developed a behavioral paradigm in which head-fixed mice use discrete tongue-reaching movements to intercept food pellets moving at constant speeds that change randomly from trial to trial. We found that mice adapted their tongue kinematics, lick onset distance, and body movement during interception training. For successful interception, mice employed a more reactive strategy early in learning, increasing tongue projection speed as pellet speed increased. Most mice developed an anticipatory strategy during late learning by initiating their licks at earlier distances for faster pellet speeds. Additionally, body movement activity ramped up as the pellet approached, indicating that mice engaged in preparatory actions several seconds before interception. Furthermore, our results show that the anticipatory strategy in our interception task partially depends on visual input, as blocking vision delayed the onset of licking and led to a shift from anticipatory to reactive tongue movement control. Similarly, visual blocking delayed the preparatory ramping of body movement. Finally, we showed that unilateral inactivation of Crus I and II impaired the anticipatory interception strategy and biased lick endpoints toward the ipsilateral site of inactivation.

### 2.1 A Novel Behavioral Paradigm for Interception

Head-fixation enables flexible integration of mouse behavior with in-vivo imaging and electrophysiology^22^. However, many head-fixed paradigms that investigate sensorimotor control are reduced to classical/operant conditioning designs or variations of them^23,24^, in which stimuli are static, response timing is loosely constrained, and successful performance does not require tight sensorimotor coupling or continuous trial-to-trial adaptation. However, these features are indispensable for ethologically relevant behavior^25–27^. To overcome these shortcomings while retaining the advantages of head fixation, we designed a behavioral paradigm that models an interception-like behavior, in which mice adaptively adjust their motor timing and kinematics in response to dynamic, continuous sensory input.

In the head-fixed mouse literature, the term “predictive” is commonly used to describe behaviors that rely on learned temporal regularities or fixed task contingencies, allowing animals to anticipate the timing of future events^28–31^. In contrast, anticipatory control in the present study refers to trial-by-trial modulation of motor onset based on an instantaneous sensory estimate of pellet speed. Although both frameworks are predictive in a broad sense, they differ fundamentally in what they predict and in the constraints they impose on motor control. Temporal expectation concerns predicting when an event is likely to occur based on prior timing statistics, whereas interceptive anticipation concerns predicting how and when to act based on the continuously evolving sensory state of a moving target. Accordingly, while blocked or repeated timing schedules are well-suited for studying feedforward motor preparation, they sometimes effectively reduce the task to a timing problem. By design, our randomly interleaved interceptive task precludes reliance on fixed temporal expectations and instead isolates predictive control in the sense of real-time state estimation guiding motor onset.

For successful interception behavior, most mice switched from using a reactive strategy in early sessions, compensating for the delay in initiating their licks by coupling their tongue projection speed to the pellet speed, to an anticipatory strategy in late sessions, where they modified the distance at which they initiated their licking to account for higher pellet speeds. These different interception strategies have been observed in human psychophysical studies when intercepting moving objects^5,32^. Our definition of these strategies reflects a more nuanced view of sensorimotor control, one that goes beyond merely categorizing responses as reactive or anticipatory based solely on response timing. It emphasizes how motor outputs are continuously tuned to dynamic sensory information, in this case, pellet speed, thereby revealing the flexible, context-dependent nature of motor planning and execution. This opens the door to investigating the neural mechanisms underlying the mapping between continuous sensory input and real-time adaptive motor responses. Additionally, both motor strategies should not be interpreted as performance-independent or equally successful; they describe the main kinematic control variables the mouse uses to successfully intercept the pellet across different speeds and sessions. Identifying these kinematic control variables will be important for future neural circuit-level investigations.

Although prior studies have successfully leveraged tongue-based reaching in mice to investigate sensorimotor control^33–36^, the behavioral paradigm introduced here is designed to address a distinct motor control problem. In sequence-licking paradigms^33,35^, the lick port moves sequentially across discrete spatial positions. Within this framework, mice are not required to estimate target velocity or generate precisely timed actions under trial-by-trial uncertainty in motion dynamics. Similarly, paradigms involving lateral displacement of the lick port^34,36^ focus on correction of tongue movements following spout contact that is displaced laterally in a discrete fixed position, rather than on motor planning under continuously varying target dynamics. Together, these approaches address complementary aspects of tongue sensorimotor control, whereas our task isolates the demands of interceptive control under trial-by-trial uncertainty in target motion.

### 2.2 Adaptive Changes in Discrete Tongue-Reaching Movements

In our interception task, mice learned to perform single, discrete tongue-reaching movements for interception. Previous protocols on forelimb-reaching reported that mice initially use their tongue for reaching and progressively shift to using their forelimb^15,16^. These observations suggest that tongue-reaching is an innate behavior in mice when the object is within reach.

While forelimb-based interception has not, to our knowledge, been studied in headfixed mice, implementing such a task variant would likely require longer training. In contrast, tongue-based interception emerged as a natural, readily learnable behavior. Importantly, our findings show that tongue-based interception exhibits some hallmark features of interception that have been well characterized in human hand-based interception (e.g., scaling of reach kinematics with target speed;^5^). This suggests that some computational principles underlying interception are similar across effector organs. Forelimb-based interception will offer a richer three-dimensional kinematic access compared to tongue-based interception, as substantial components of tongue motion occur intra-orally and are therefore not directly observable^37^. Extending this paradigm to forepaw-based interception represents an important direction for future work.

Furthermore, the discrete tongue movements observed in this study are characterized by a highly adaptable motor motif, comprising distinct projection, probing, and retraction phases. This motif can be reiterated at intervals as brief as 15 milliseconds before full tongue retraction, generating a complex lick trajectory indicative of real-time, online corrective adjustments. These observations align with prior reports describing different phases of tongue movement during stereotyped rhythmic water-licking behaviors^36,38^. However, in contrast to these earlier studies, our findings reveal that this tongue-specific motor motif exhibits significant plasticity, adapting dynamically to variations in pellet speed and the stage of learning. These findings lay the groundwork for investigating the neural mechanisms of discrete versus rhythmic tongue movement^39^, alongside those driving adaptive tongue kinematics.

### 2.3 Learning-Dependent Changes in Body and Whisker Movement

Mice performing this task exhibited adaptive changes in body and whisker movements. As learning progressed, the onset of ramping of both body and whisker movements shifted earlier. The amplitude of these movements increased in a distance-dependent manner, and body movement peaked earlier, then decreased as whisker movement began to rise, possibly allowing for a more stable posture before tongue movement^40^. These preparatory whisker and whole-body movements are similar to the anticipatory postural adjustments that occur before movement and correlate with improvements in reaching performance in humans and cats^40–42^. The distance dependence of these preparatory movements suggests that they reflect tracking of the sensory environment^43,44^, specifically, the position of the pellet, which may aid in guiding interceptive actions.

### 2.4 Partial Reliance on Monocular Cues

Visual blocking significantly reduced the general hit rate for pellet speeds above 4 cm/s, indicating that visual information is particularly important for intercepting fast-moving targets. In contrast, blocking vision led to a comparable reduction in the single-lick hit rate across all pellet speeds, reflecting a general decrease in the accuracy of individual tongue reaching movements rather than a speed-specific failure of interception. This reduction in single-lick accuracy at low pellet speeds does not imply that vision is strictly required under these conditions, but rather that visual input allows for fine calibration even when slower targets are possibly being tracked using additional sensory modalities, such as the whiskers or olfaction.^45,46^.

The fact that monocular blocking impaired the animal’s ability to intercept fast-moving pellets shows that mice can use monocular cues (e.g., looming) for our interception task; this is similar to mice using monocular cues (e.g., motion parallax) for distance estimation when jumping over a gap between two platforms^47^. After visual blocking, mice initiated licks earlier at slow pellet speeds but later at high pellet speeds. Visual blocking did not affect kinematics, except for tongue projection speed, which correlated positively with pellet speed. These modulations in lick onset distance and lick projection speed reflected a shift in interception strategy, becoming more reactive and less anticipatory. In addition, visual blocking affected preparatory whisker and whole-body movements, delaying the onset of body and whisker ramping and shifting the timing coordination pattern. Future work may investigate neural mechanisms for computing different optical variables (e.g., visual angle and angular velocity) that aid anticipatory interception^48–50^.

### 2.5 Lateral Cerebellar Circuits Are Necessary for Task Execution

The involvement of the lateral cerebellum in interception could either be related to sensory acquisition^20,51–54^ or controlling tongue-related motor kinematics.^55–60^. Discerning how these different contributions are mapped and organized within cerebellar circuits will require recording neural activity during our task. Our pharmacological inhibition approach suggests that cerebellar Crus I and II contribute to learned motor kinematics of tongue movement. After inhibition, the tongue projection speed increased regardless of the pellet speed. Additionally, there was a significant deflection in the lateral positioning of the tongue. This result replicates recent findings showing that optogenetic stimulation biases the tongue lick angle toward the ipsilateral side of the manipulation^60^. Moreover, there was a significant shift in lick onset distance, impairing the ability of mice to time their movement adaptively across different pellet speeds. Future studies should investigate cerebellar cell-type-specific contributions to the various perceptual and motor aspects of this task.

Tongue motor control is implemented through coordinated activity across bilateral hypoglossal motor nuclei, receiving direct and indirect inputs from deep cerebellar nuclei^61^, suggesting that effective interceptive motor control of the tongue likely depends on bilateral cerebellar contributions rather than a strictly unilateral mechanism. Additionally, sensory information (e.g., visual and whisker) guiding interceptive action can reach both cerebellar hemispheres via a combination of cerebro-cerebellar projections and subcortical brainstem pathways. Together, these anatomical considerations suggest that both cerebellar hemispheres contribute to task performance. Systematic investigation of cerebellar laterality, as well as perturbations of additional cortical and subcortical regions such as the posterior parietal cortex, represents important directions for future work.

## 3 Material and Methods

### 3.1 Animals

Behavioral experiments were performed on fifteen adult male C57BL/6 mice and heterozygous Pcp2::Cre mice (B6.Cg-Tg [Pcp2Cre] 3555jdhu/J; Jax stock #010536) aged between 12-16 weeks. Before the start of behavioral experiments, mice were group-housed (2 mice per cage) and held under a controlled 12h/12h light/dark cycle with adlibitum access to food and water. All animal experiments followed guidelines approved by the Institutional Animal Care and Use Committee in an Association for Assessment and Accreditation of Laboratory Animal Care (AAALAC International)-accredited facility.

### 3.2 Surgical Procedure

All mice underwent stereotactic surgery to expose the left lateral cerebellar lobules Crus I and II, and to fix a clear glass window and a metal head plate on the skull. This allowed for head fixation and simultaneous optical access to these lobules, as described in previous reports^62,63^.

Briefly, before surgery, a small silicon access port was drilled into a 5 mm coverslip (170 *µ*m thick). Mice were anesthetized with a mixture of Medetomidine (0.3 mg/kg), Midazolam (4 mg/kg), and Butorphanol (5 mg/kg) in 0.9% saline solution (intraperitoneal injection). Eye ointment was applied to protect the eyes from drying during the surgery. After shaving the hair and sterilizing the skin with Iodine solution, a 2% lidocaine solution was applied to the sterilized region, and an incision was made to remove the skin.

Once the skull was exposed, muscular attachments covering the left cerebellar hemisphere were removed carefully. The posterior suture line was used as an anatomical landmark, and a 5 mm diameter skull flap was drilled using a dental drill (EXL-M40, Osaka). The central point of the skull flap was 2.5-3.5 mm lateral and 1 mm posterior from the midpoint of the posterior suture line. The coverslip with the silicon access port was firmly pressed into place over the exposed cerebellar area, and a drop of superglue was applied to the perimeter of the coverslip. Dental adhesive resin cement (Super-Bond C&B) was used to attach a metal headplate over the coverslip.

After surgery, mice were placed in their home cage on a heated plate until they regained body mobility. Carprofen (5 *µ*g/g body weight, intraperitoneal) and Buprenorphine (0.1 *µ*g/g body weight, subcutaneous) were given to reduce postoperative inflammation and pain, respectively. Mice were given a recovery period of at least 5 days before handling began. All behavioral experiments were done during the dark period of the 12/12 hour light/dark cycle.

### 3.3 Behavioral Training and Testing

#### 3.3.1 Habituation

After the recovery period, mice were handled for 30 minutes each day for 5 days. A modified 3D-printed housing^64^, similar to that in the experimental setup, was used to handle and habituate mice, facilitating their voluntary entry and 1-3 minutes of stay.

#### 3.3.2 Food and Water Deprivation

Food and water deprivation aimed to encourage mice to use their tongue to intercept the moving pellet. Mice were started on a water-deprivation schedule, as shown in our earlier pilot experiments, to encourage them to extend their tongues under head fixation. Each mouse received 2 g of water gel daily in their home cage. Each day, water-deprived mice were head-fixed and habituated to the setup for 10 to 30 minutes. During this time, mice could consume up to 1 ml of water from a spout located 2-3 mm away using their tongue. Mice started licking within 3-4 days. Once mice readily licked from the spout, water deprivation was discontinued, and mice were allowed ad libitum access to food and water for 1 day. Once mice regained their original weight, food deprivation started. Mice were kept between 85 to 90% of their original body weight. During the first 4 days of food deprivation, mice were head-fixed on the setup and presented with 10 mg food pellets (Purified Rodent Tablets, TestDiet) on a stationary platform positioned 2.5-3 mm underneath their mouth. Once mice readily extended their tongue to obtain the food pellet, behavioral testing and data acquisition began.

#### 3.3.3 Behavioral Testing

To habituate mice to the moving platform, mice first completed a single session during which the platform moved at the lowest two pellet speeds (1-2 cm/s). After this session, mice completed six training sessions to optimize their behavior for intercepting pellets moving at constant speeds from 1 to 7 cm/s, which were randomly selected from trial to trial. This randomization was important to prevent mice from relying on fixed temporal expectations. Each training session lasted between 1 and 1.5 hours. All sessions were conducted under blue light illumination. At the beginning of each session, mice were guided into a modified 3D-printed body-restraint tube^64^ and head-fixed. The right edge of the platform holding the pellet was aligned to the midline of the mice and moved down 3-4 mm relative to the snout. The platform’s position was kept the same across all training sessions. Following alignment, the platform was moved 25-27 cm to the beginning of the guiding linear rail.

Each session comprised 70 trials, where pellet speeds between 1 and 7 cm/s were counterbalanced and randomly assigned at each trial. A random inter-trial interval between 15 and 25 seconds was added, followed by automatic pellet-refilling if necessary. Each trial started with a 500 Hz tone from a buzzer located on the moving platform, which played until the platform reached a distance of 2.5 cm beyond the midpoint of the mice (defined as the middle point of the lower jaw during rest). White noise at 60 dB was played throughout each session to mask mechanical noise from the platform’s movement along the rail. At the end of each session, mice were returned to their home cage and provided with complementary food pellets to ensure their daily intake was 2.5-3 g. If the animal’s weight dropped below 85%, the amount of food pellets provided at the end of each session was increased. If the weight dropped below 80%, animals were removed from the experiment.

#### 3.3.4 Visual Blocking Experiments

Four mice were trained for up to six days on the regular interception task. Their performance was then recorded without or with a barrier blocking the view of the approaching food pellet. A 4 x 4 cm piece of black aluminum foil (Cinefoil) was placed near the left eye to block visual input from the left side (the same side where the pellet comes).

#### 3.3.5 Intra-Cerebellar Muscimol Injections

Four mice that underwent interception training were used to test the involvement of lateral cerebellar circuits. Muscimol (Tocris), a potent GABAA agonist, was injected unilaterally, on the left side (the same side from which the pellet approached), 150 µm from the cerebellar cortical surface through the silicon port of the coverslip. A quartz pipette was pulled using a laser puller (P-2000, Sutter Instrument) and beveled at 35 degrees (BV-10, Sutter Instrument) to obtain a pipette tip with a 10–15 µm opening diameter. The pipette was then backfilled with 1 ng/nl muscimol solution. 120-140 nl of muscimol were injected through the silicon port in Crus I near the border with Crus II. After waiting 15-20 minutes, the session began. In the following session, a matching volume of saline was injected into the same brain region as a control.

### 3.4 Behavioral Experiment Hardware: Design and Setup

The hardware shown in Figure 1A comprises three primary components:

1. A linear belt actuator-rail and motor unit. A linear belt actuator (W45-15 linear module, CCM) was combined with a NEMA 23 servo motor (CPM-SDSK-2310S-RLN, Clearpath Teknic). This motor provided an integrated high-precision positional and velocity control solution, along with a silent operation mode and positional feedback router. Together, these components allowed us to increase pellet speed to 20 cm/s with minimal mechanical noise and to support the weight of a 400 g micromanipulator (MM3333, Narishige) to which the pellet platform was attached. An Arduino wrapper was used to access the ClearCore C++ Motion and I/O Library, enabling us to control the motor’s position and speed according to the task design. The linear belt actuator and the micromanipulator holding the pellet platform were mounted on a breadboard connected directly to an air table. They were mechanically separated from the mouse housing stage, which was positioned under the microscope. This design helped to ensure that any vibration produced by the motor was dissipated by the air table before reaching the mouse stage.
2. An automatic pellet-refilling mechanism. Pellet-refilling between trials was fully automated. The pellet platform was 3D-designed and printed (Figure 1A, sub-panel). The design allowed the pellet to rest on a concave platform surface; this concave surface was connected to the platform’s bottom with a small light path, allowing laser light (650 nm transmitter, KY008 Arduino module) to pass. If a pellet was present, the laser light was blocked. If no pellet was present, a photosensitive light sensor (photosensitive Light Dependent Resistor (LDR) ST0250 Arduino module, sunFounder) detected the incoming light, which triggered the release of a pellet from a pellet dispenser (PD-020D, O’Hara & C., Ltd). The pellet was delivered through a transparent silicone tube to the pellet platform.
3. Two high-speed cameras Two monochrome cameras (BFS-U3-04S2M-CS USB3.1 Blackfly, Teledyne FLIR) were positioned in orthogonal planes to record mice from a bottom and side view, which allowed us to determine the 3D coordinates of tongue movements. The cameras had low distortion, variable focal length, and high-resolution lenses (4.4-11mm FL Varifocal Lens, Edmund optics). An Arduino microcontroller provided triggers to synchronize both cameras using GPIO camera cables. Spinnaker software (Spinnaker-2.7.0.128 for Windows) was used for video acquisition. Four infrared LEDs (M940L3, M680L5; Thorlabs Inc.) were used to illuminate the mouse from both the bottom and side views. One blue LED (M470L5; Thorlabs Inc.) illuminated the experimental setup. All experiments were done in the same lighting conditions.

These three components were securely mounted on two aluminum breadboards (Thorlabs Inc.), each measuring 300 mm x 900 mm x 12.7 mm. The system was managed via an Arduino microcontroller (Mega 2560 ATmega2560, Arduino) programmed with custom scripts to control and synchronize all hardware functions.

### 3.5 Behavioral Recording, Acquisition, and Processing

Synchronized bottom and side-view camera recordings were acquired at 501 Hz. Each trial video was saved in .avi format. DeepLabCut^17^ was used to track specific body parts. Forty videos from eight mice in different conditions and stages of learning were used to extract frames for training a resnet50 model. We extracted frames manually since the tongue appears only briefly during each trial. From the side view, we labeled the tip of the snout, the left and right paws, the pellet, and the tip and base of the tongue. From the bottom view, we labeled the left and right paws, the midpoint of the lower jaw, the tip and base of the tongue, and the right edge of the platform. In total, we labeled 5000 frames and trained two separate side- and bottom-view networks. We ran the networks on 20 held-out videos to confirm accurate tracking and manually checked performance. If network performance was not adequate, we retrained the network. Usually, we only needed one retraining step. In total, 500,000 training iterations were needed for the network to achieve a 1.5-pixel error in the network evaluation. The trained network was used to analyze all videos via the OIST high-performance GPU cluster. All tracking data was saved in Excel format. A MATLAB (MathWorks, Inc.) pipeline with the following three modules was written to process and analyze all behavioral tracking data:

1. Tongue manual tracking module The main aim of this routine was to extract 3D tongue kinematics similar to the work by Bollu et al. 2021^36^. Images from the bottom and side views were used to track X, Y, and Z coordinates of the tongue tip. X and Y coordinates (anteroposterior and dorsoventral axes) were tracked from the side view, while the Z coordinate (mediolateral axis) was tracked from the bottom view. The DeepLabCut tongue tip tracking from both camera views was used as the initial detection layer. The frames detected as lick active were processed into another manual layer. Using an interactive user interface, we corrected the start and end times for each lick and, if necessary, annotated the X, Y, and Z coordinates of the tongue tip. After the X, Y, and Z coordinates were manually checked on both side and bottom views, they were smoothed using a cubic smoothing spline (csaps, MATLAB) with a smoothing parameter p=0.1. A common initial starting point was subtracted from all X, Y, and Z coordinates, and the resulting pixel distances were converted to millimeter scale using the calibration indices for the bottom and side view cameras. Figure 1C shows an example of tracking the tongue tip during a single lick from both the bottom and side view cameras. Positional measurement error was estimated from the discrepancy in the total three-dimensional tongue-tip path length between manual and automated tracking (approximately 0.5 mm). This value was used as a conservative threshold for endpoint position analyses. Different tongue phases were segmented similarly to previous work by Bollu et al.^36^, although with slight differences. Segmentation for each lick was performed along the dorsoventral X and Y axes. The projection phase was defined as the time between the beginning of the lick and before the first local minimum. The retraction phase was defined as the first local minimum after inverting the trace. The probing phase was defined as the period between the projection and retraction phases. For the complex licks, the number of peaks that exceeded half of the first projection phase amplitude was quantified using the “findpeaks” function in Matlab, and then used to segment the single licks into smaller simple lick segments with the three phases of movement. After applying this automatic algorithm, a user interface allowed the user to confirm the selection of all different inflection points between phases. Additionally, a long short-term memory (LSTM) deep network for time series segmentation was trained on several manually segmented traces and then used as a complementary, optional tool for automatically segmenting the traces. Tongue instantaneous speed was calculated as the Euclidean distance (in millimeters) between consecutive 3D trajectory points, divided by the time interval (in seconds) between those points. The lick onset distance was defined as the relative distance of the pellet from the animal’s midline at the start of the lick. Lick duration was defined as the total time of the lick. The lick speed parameter was also defined for each phase of the lick, as described above. The lick endpoint was defined as the 3D coordinates of the tongue tip at the onset of the retraction phase, since this point marks a clear end of the lick-reaching attempt. The lick angle was defined as the angle between a line connecting the tongue tip and lower jaw to a hypothetical vertical midline crossing the lower jaw and snout. Figure S1 shows segmentation examples of two distinctive types of discrete licks that mice made during the experiments. Rhythmic licking is defined as a full tongue movement, with all three kinematic phases, repeated more than once in the trial, with the explicit condition that the tongue is fully retracted between movements. This is in opposition to discrete single licks, which are defined as a single tongue reaching movement that can be simple or complex, with no preceding or consecutive licks in any part of the trial.
2. Motion energy of body parts module The main aim of this module was to quantify the movement of body parts not amenable to point-based tracking in our setup (the whisker pad and visible part of the whole body) using instantaneous motion energy, defined as the mean intensity of the pixel difference between consecutive frames. A user interface was created to allow us to select multiple regions of interest, which were used to calculate the average difference between frames. All motion energy traces were normalized and smoothed using a moving average filter with a window size of 100 milliseconds. Traces are expressed in arbitrary units of motion energy. Figure 1B shows examples of the regions-of-interest for the whole-body and whisker pad, and the resulting motion energy traces.
3. Pellet and distance tracking module Pellet tracking data extracted from the side view camera was used to define hit (animal obtaining the pellet and raising it more than 2 mm), miss (animal licking but failing to contact the pellet), or no-lick trials (animal not licking at all). An automatic algorithm was developed to determine each trial’s success based on the existence of the pellet on the platform before and after each lick. If the pellet moved more than 2 mm towards the mouse in the ventro-dorsal direction, it was considered a successful lick. Another user interface was also developed to gather manual feedback and ensure the accuracy of the automatic detection. As the pellet moved at a constant speed, the distance the object moved in each frame was derived and normalized to the midpoint of the animal, defined as the point at which the platform’s right edge aligned with the animal’s midline.

### 3.6 Behavioral Data Statistical Analysis

All statistical analyses were performed using MATLAB software (R2022b, MathWorks) or GraphPad Prism software (Prism 10.4.0). All statistical tests were two-tailed, assigning significance if the p-value was less than 0.05. All data presented in the text and figures reflect the mean ± SEM. Significance levels in the graphs were determined based on the following threshold : * p *<* 0.05; ** p *<* 0.01; *** p *<* 0.001; **** p *<* 0.0001 unless otherwise stated. If there were multiple comparisons, p-values were corrected using the Benjamini-Hochberg method^65^. Fixed-effect results from either the two-way analysis of variance (ANOVA) or linear mixed-effects models were reported in the figures, with significance levels indicated. For correlations, the two-tailed Pearson correlation was used after removing identified outliers using Robust regression and Outlier removal (ROUT) in Graphpad Prism software. For circular data, a two-way ANOVA for circular data with interactions was used from the Matlab circular analysis toolbox^66^

#### 3.6.1 Logistic linear mixed-effects model

To identify behavioral factors underlying the binary outcome of interception success, we employed a logistic linear mixed-effects model. This model included both fixed effects, which capture population-level trends across all animals, and random effects, which account for individual differences between mice. We used MATLAB’s “fitglme” function with a logit link and a binomial distribution to model success, and fitted the model using the Laplace approximation for maximum likelihood estimation. Fixed effects included the main effects and all two- and three-way interactions between learning stage, lick variables (lick duration, lick onset distance, and lick projection speed), and pellet speed. The inclusion of three-way interactions allowed us to assess how learning modulates the interaction between different lick variables and different pellet speeds, possibly revealing specific behavioral strategy changes. To account for inter-individual variability in baseline performance and sensitivity to pellet speed, we included a random intercept and a random slope for pellet speed for each mouse.

where:

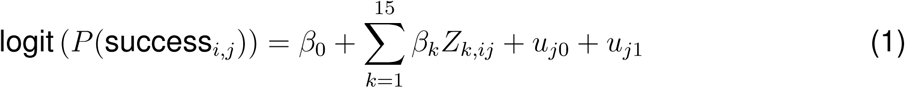

- *P* (success*_i,j_*): The probability of success for trial *i* and subject *j*.
- logit(*P*): The log-odds transformation of the probability *P*, defined as log *<u>^P^</u>*.
- *β*_0_: The fixed-effect intercept, representing the overall baseline log-odds of success.
- *β_k_*: The fixed-effect coefficient for predictor *Z_k_*, representing the population-level effect of the *k*-th predictor.
- *Z_k_*: The *k*-th fixed effect term, representing all main effects and interaction terms involving learning stage, lick duration, lick onset distance, lick projection speed, and pellet speed.
- *u_j_*_0_: The random-effect intercept for subject *j*, capturing mouse-specific deviations from the overall intercept.
- *u_j_*_1_: The random-slope for pellet speed in subject *j*, allowing for the effect of pellet speed to vary across mice.

#### 3.6.2 Strategy Modulation Index Calculation

To quantify the extent to which animals employed an anticipatory versus reactive strategy during interception, we defined a strategy modulation index (MI) based on features of licking behavior that influenced interception success as per the logistic linear mixed-effects model, including lick onset distance and its interaction with pellet speed, as well as the interaction between tongue projection speed and pellet speed. Specifically, three key features were used in the calculation of the index: (1) the median lick onset distance across all pellet speeds, (2) the correlation between lick onset distance and pellet speed, and (3) the correlation between tongue projection speed and pellet speed. The score quantifies the directional modulation of these features. A higher score (closer to 1) indicates a more anticipatory strategy where mice exhibit more negative lick onset distances (generally licking earlier regardless of the pellet speed), a negative correlation between lick onset distance and pellet speed (licking earlier for higher pellet speed), and a near-zero correlation between tongue projection speed and pellet speed. Conversely, a lower MI score (closer to 0) indicates a more reactive strategy with more positive lick onset distances (delayed lick initiation), and positive correlations between lick onset distance and pellet speed, as well as tongue projection speed and pellet speed. Each of the three features was normalized to a range between 0 and 1, and then the MI was calculated as follows:

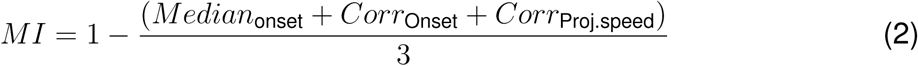

#### 3.6.3 Clustering Analysis

To identify interceptive strategies, we applied k-means clustering in MATLAB using the “kmeans” function to the MI scores computed for each mouse in each session. Each session was treated as an independent data point characterized by a single MI value. The number of clusters was fixed at k=2, reflecting the hypothesis that animals express one of two dominant strategies at each stage of learning. Cluster centroids were used to assign strategy labels.

## Supplementary Materials

**Supplementary Figure 1:**
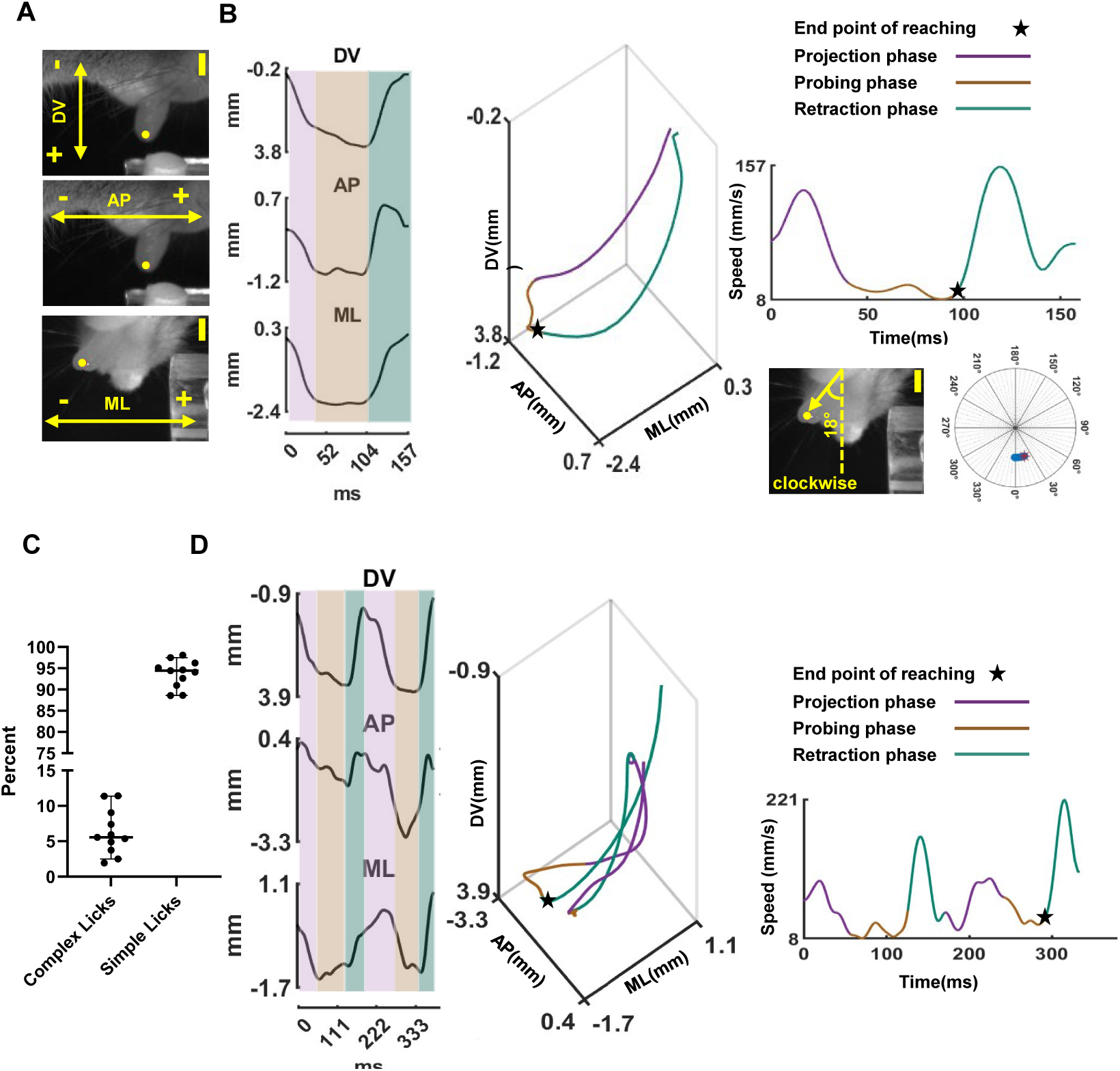
Single and complex lick segmentation. (A) The first two rows show the side camera view, which was used to extract the anteroposterior (AP) and dorso-ventral (DV) axes. The third row shows the bottom-view camera used to extract the medio-lateral (ML) axis. Scale bar = 3 mm. (B) Left: Segmentation of lick phases in all three axes for a representative simple lick. Colors represent the different motor phases (projection, probing, and retraction). Middle: 3D reconstruction of the tongue tip trajectory. Right: The tongue speed profile segmented into phases (upper panel) and the maximum tongue angle at the reach endpoint (lower panel). The angle is measured clockwise in degrees. (C) Percentage of simple versus complex licks. The error bars denote the SEM (N = 11 mice). (D) Lick segmentation, lick trajectory, and lick speed profile for a representative complex lick.

**Supplementary Figure 2:**
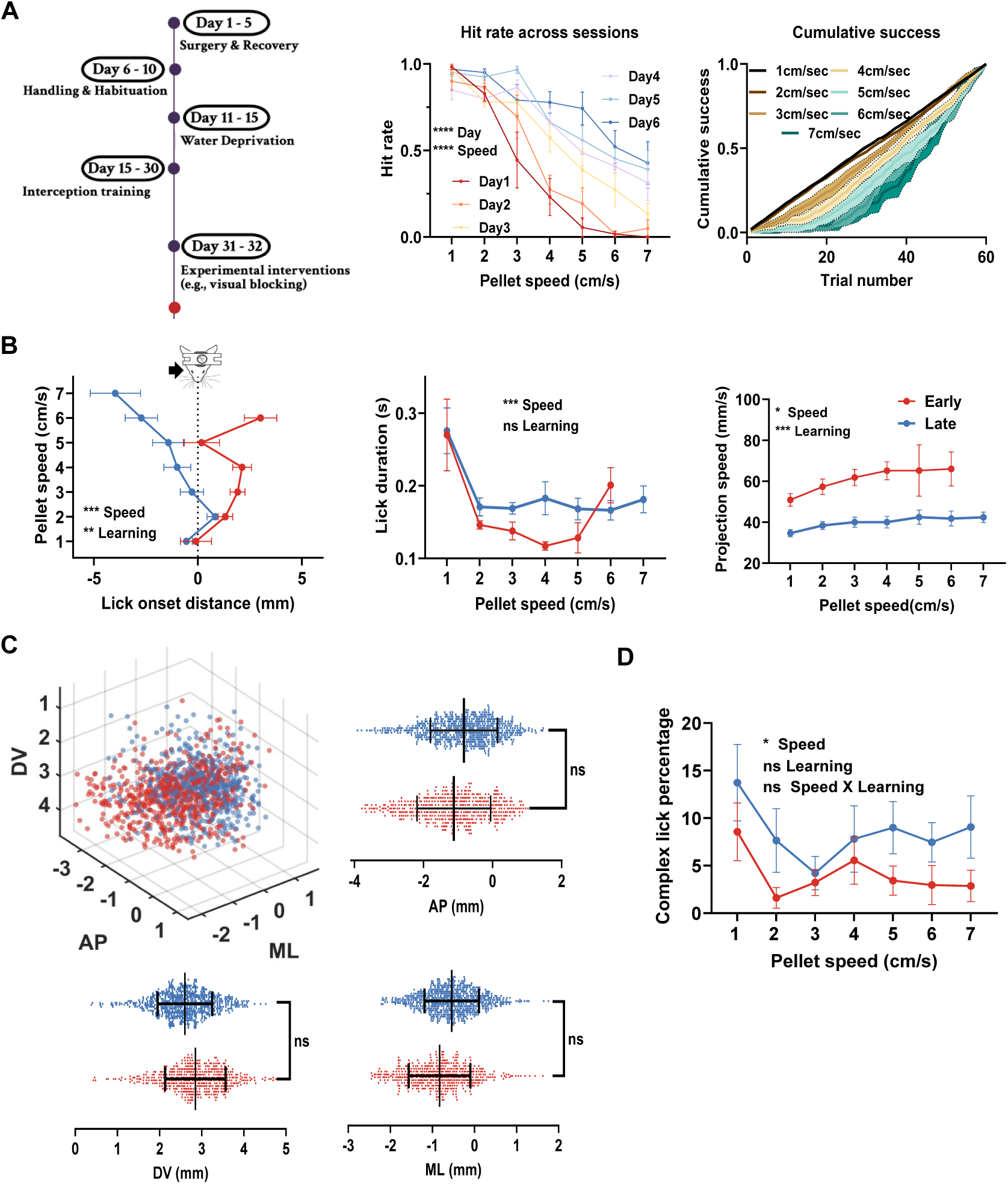
Gradual motor learning, kinematics of successful single licks and spatial distribution of lick endpoints. (A) Left: A schematic of the experimental timeline. Middle: Hit rate versus pellet for different days across six mice. Right: Cumulative success versus the total number of trials over six training days for different pellet speeds. (B) Left: Lick onset distance of successful single licks in early and late sessions. Middle: Lick duration in successful single licks. Right: Lick projection speed in successful single licks. (C) The spatial distribution of the first lick endpoint in early and late sessions (each dot is a single-lick in each mouse). AP, DV, and ML indicate the anteroposterior, dorso-ventral, and mediolateral axes, respectively. (D) Complex lick percentage as a function of pellet speed. The complex lick percentage shows a non-significant decrease across sessions and a significant increase at a pellet speed of 1 cm/s. All data are presented as mean ± SEM, except for panel C, which is shown as mean ± standard deviation to illustrate variability in lick endpoint positions. Analyses were performed using linear mixed-effects models or two-way repeated-measures ANOVAs. Significance levels of the main and interaction effects are reported as *p*<*0.05, **p*<*0.01, ***p*<*0.001, and ****p*<*0.0001

**Supplementary Figure 3:**
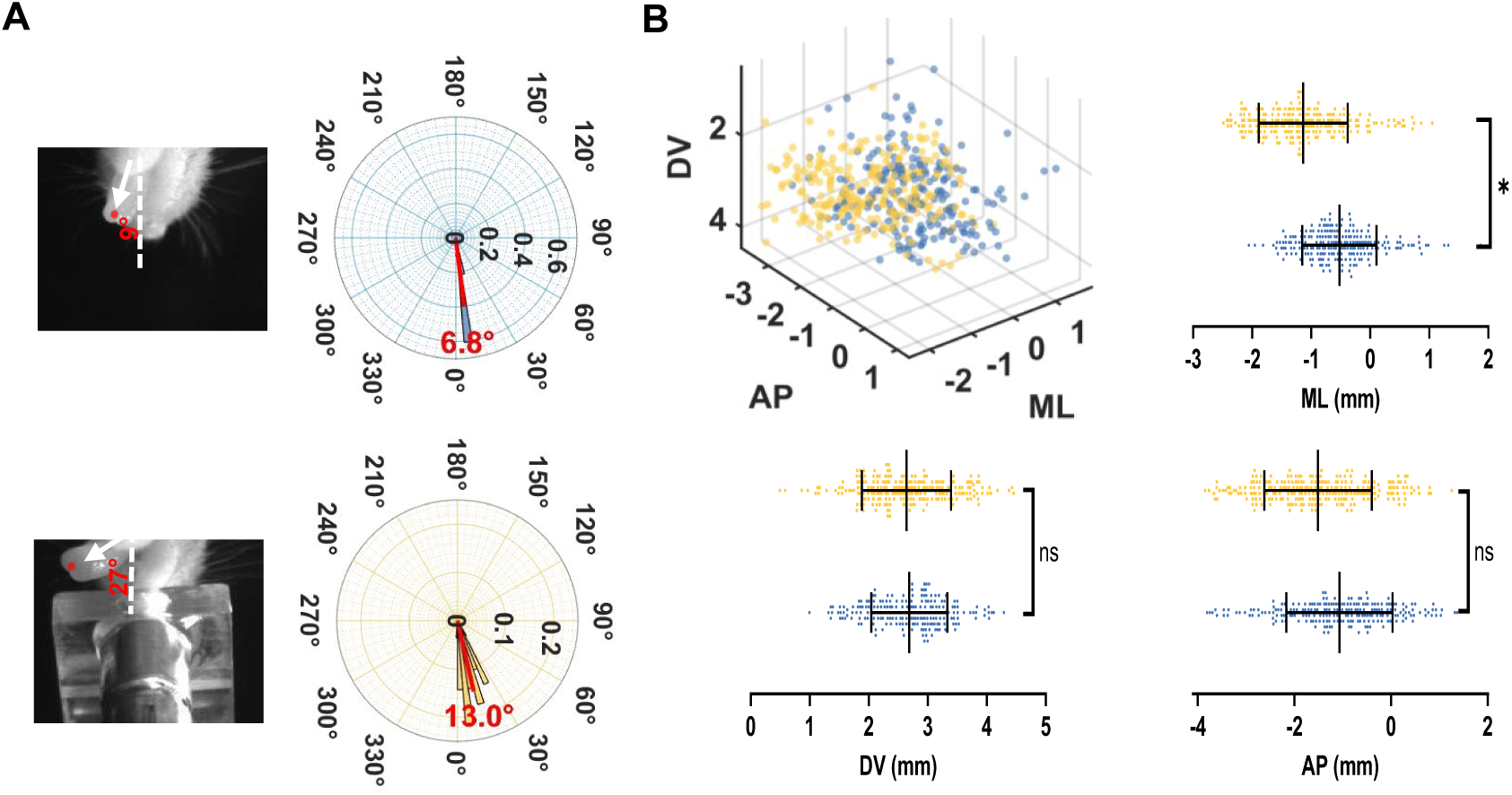
Effect of muscimol on the spatial distribution of lick endpoints. (A) Upper: An example of the maximum reach endpoint angle (left) and the angle distribution across different trials. The mean angle is denoted in red. Lower: Same as the upper panel, but now after muscimol injection. There was a significant increase in the endpoint angle *F* (1, 42) = 10.854*, p* = 0.002. Analyses were conducted using the circular statistics toolbox^66^. (B) Spatial position of the reach endpoint of all licks in all axes. Only the mediolateral axis showed a significant difference after muscimol injection.

**Supplementary Figure 4:**
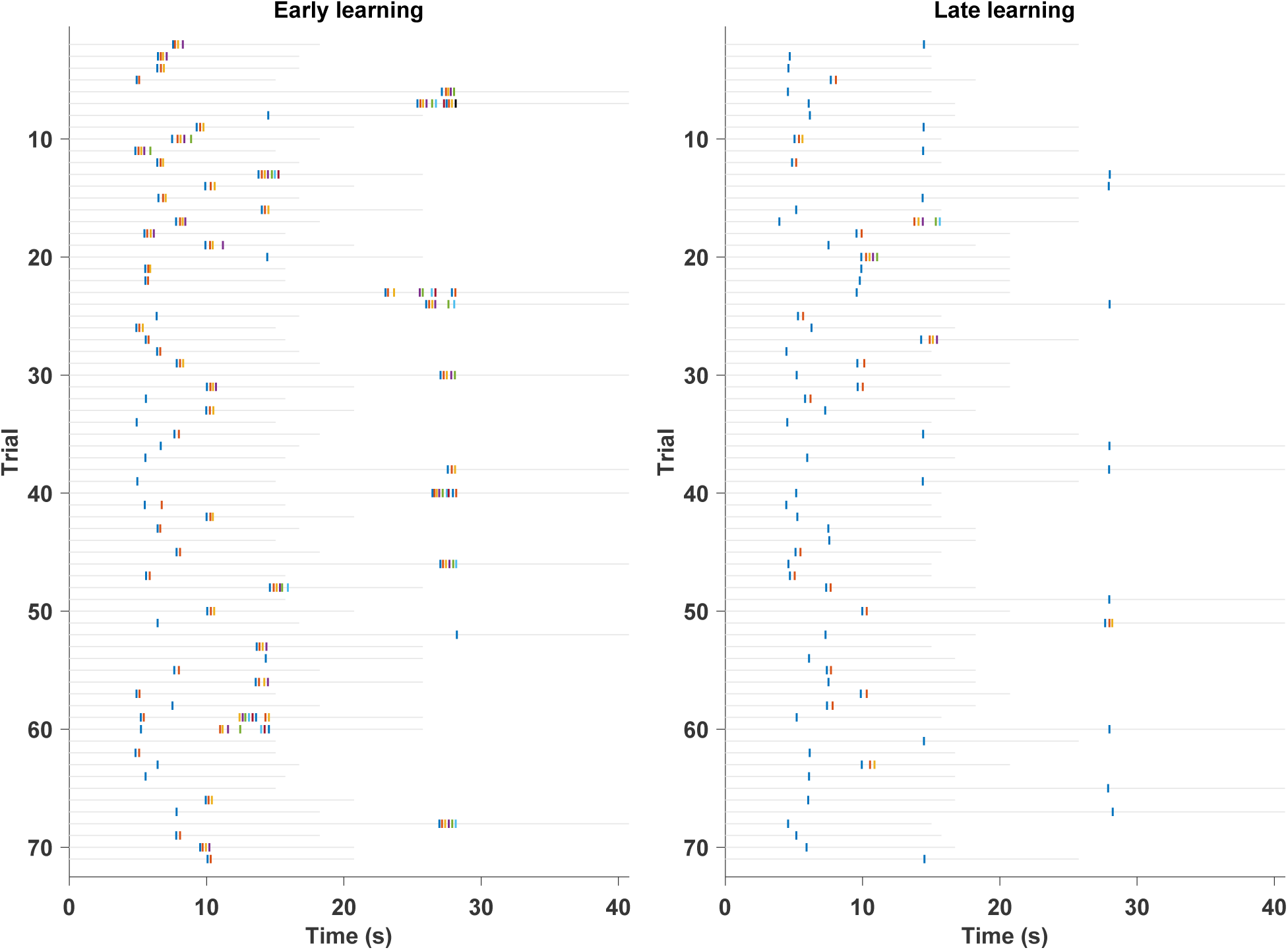
Raster plot showing the progression of licking and emergence of single licks in an example mouse. Left: Early session. Each gray horizontal line indicates the duration of the trial from the beginning of pellet movement to the end. The vertical colored lines indicate the start of individual licks in each trial. Licks are colored differently for visualization. Right: Same as the left panel, but for the late session.

**Supplementary Table 1:**
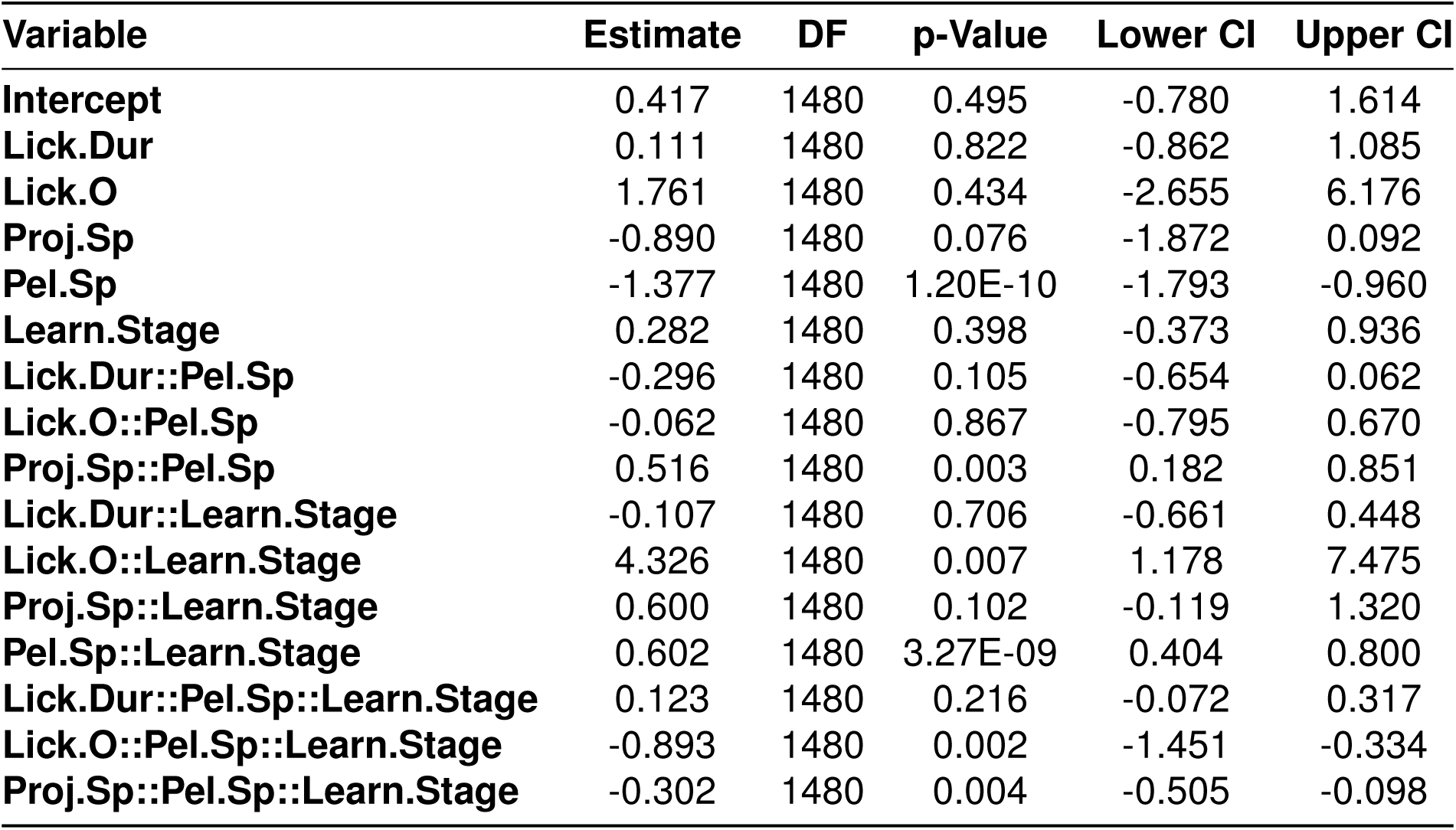
Summary of fixed effects from the logistic linear mixed-effects model predicting interception success. Significant effects indicate behavioral or task variables influencing performance across learning and pellet speeds. Abbreviations: Lick onset distance (Lick O), lick duration (Lick Dur), Lick projection speed (Proj.Sp), Pellet speed (Pel.Sp), Learning stage (Learn.Stage).

## 3.7 Resource availability

### 3.7.1 Lead contact

Requests for further information and resources should be directed to and will be fulfilled by any of the corresponding authors.

## Data and code availability

The lead contacts will share all data reported in this paper upon request. Additionally, all datasets and analysis software used for tracking the tongue will be deposited online upon publication.

## Supplemental information

There are four supplementary figures, one supplementary table showing the results of the logistic linear mixed effect model in detail in early and late learning, and two supplementary movies. Supplementary Movie 1 shows different types of licks (simple short, simple long, and complex lick); Supplementary Movie 2 demonstrates the change of lick onset distance and projection speed of the tongue late in learning while intercepting a pellet at high pellet speed (7 cm/s).

## Acknowledgments

The authors thank the OIST Animal Resources Section, Imaging Section, Kazuo Mori, and Sigita Augustinaite for their help and technical support, Pavel Puchekov in the OIST High-Performance Computing Section for the setup illustration in Figure 1, and the OIST Graduate University for generous financial support. This work was supported by the JSPS Fellowship/KAKENHI Grant (22J10169) awarded to M. ELT. and KAKENHI Grant (23H02454) awarded to B.K.

## Author contributions

Conceptualization, M.ELT, C.G. and B.K; methodology, M.ELT and C.G..; investigation, M.ELT and C.G.; writing – original draft, M.ELT, C.G.; writing – review & editing, M.ELT, C.G. and B.K; funding acquisition, M.ELT and B.K; resources, M.ELT and B.K; supervision, B.K.

## Declaration of generative AI and AI-assisted technologies

While preparing this manuscript, the authors used ChatGPT/Grok to paraphrase and correct grammatical and punctuation mistakes. After using this tool, the authors reviewed and edited the content as needed and take full responsibility for the publication’s content.

